# How Communication Pathways Bridge Local and Global Conformations in an IgG4 Antibody: a Molecular Dynamics Study

**DOI:** 10.1101/2021.06.23.449604

**Authors:** Thomas Tarenzi, Marta Rigoli, Raffaello Potestio

**Affiliations:** Department of Physics, University of Trento, via Sommarive 14, I-38123 Trento, Italy; INFN-TIFPA, Trento Institute for Fundamental Physics and Applications, I-38123 Trento, Italy

**Author notes:** Contributed equally to this work (Thomas Tarenzi, Marta Rigoli).

## Abstract

The affinity of an antibody for its antigen is primarily determined by the specific sequence and structural arrangement of the complementarity-determining regions (CDRs). Recently, however, evidence has accumulated that points toward a nontrivial relation between the CDR and distal sites on the antibody structure: variations in the binding strengths have been observed upon mutating amino acids separated from the paratope by several nanometers, thus suggesting the existence of a communication network within antibodies whose extension and relevance might be deeper than insofar expected. In this work, we test this hypothesis by means of molecular dynamics (MD) simulations of the IgG4 monoclonal antibody pembrolizumab, an approved drug that targets the programmed cell death protein 1 (PD-1). The molecule is simulated in both the apo and holo states, totalling 4*μs* of MD trajectory. The analysis of these simulations shows that the bound antibody explores a restricted range of conformations with respect to the apo one, and that the global conformation of the molecule correlates with that of the CDR; a pivotal role in this relationship is played by the relatively short hinge, which mechanically couples Fab and Fc domains. These results support the hypothesis that pembrolizumab behaves as a complex machinery, with a multi-scale hierarchy of global and local conformational changes that communicate with one another. The analysis pipeline developed in this work is general, and it can help shed further light on the mechanistic aspects of antibody function.

**Synopsis:** Antigen binding restricts the conformational variability of the therapeutic antibody pembrolizumab in an interplay between the paratope and hinge region, mediated by a full-scale interaction network.

**Graphical TOC Entry:** 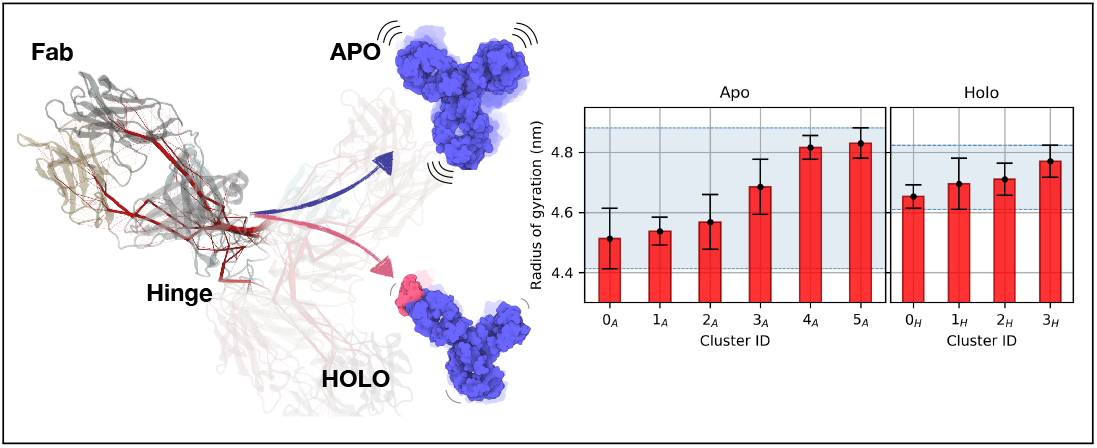

## Introduction

The number of monoclonal antibodies (mAbs) employed for therapeutic applications dramatically increased in the recent years: from 1997 to 2013, 34 mAbs-based pharmaceuticals were approved in US or EU, while from 2014 to 2020, in only 7 years, the number of approved mAbs was 61.^1^ mAbs have been developed to treat a large variety of conditions, including cancer, autoimmune diseases and, very recently, COVID-19.^2,3^

Even in the case of naked immunoglobuline drugs, which do not involve conjugation with radionuclides or small molecules, ^4,5^ engineering of the antibody sequence is routinely performed to optimize its therapeutic efficacy for a given function, through successive steps of humanization, affinity maturation, and modifications aimed at overcoming challenges in stability and manifacturing.^6^ On the one side, the selection of the isotype, and therefore those structural/dynamical features of the constant region that come with it, leads to different immune responses, and is thus performed on the basis of the planned application;^7,8^ on the other side, modifications of single residues can determine a higher therapeutic efficacy, as in the case of those mutations introduced in the Fc domain to enhance effector function and recruitment of additional proteins.^9–11^ A remarkable example is the single-residue mutation that, in the hinge of IgG4 antibodies, prevents Fab-arm exchange.^12–14^

Modifications of this type, which may be distributed throughout the whole antibody sequence, are usually introduced for reasons that are not directly linked to antigen affinity modulation. Engineering efforts intended to increase specificity and affinity are in fact mostly focused on the residues of the six loops comprising the complementarity-determining region (CDR), because of their preeminent role in antigen binding.^15^ However, the CDR loops are not the only possible loci of intervention; it is experimentally shown that both mutations near *and* far from the antigen-binding site can affect affinity.^16^ Such mutations act by modulating the interdomain conformational dynamics of the antigen-binding fragment, which eventually reflects on the paratope, namely the antigen binding site. On a similar note, NMR relaxation dispersion experiments allowed researchers to detect important fluctuating residues that are not located in the CDR, whose point mutation can nonetheless increase antigen-antibody affinity.^17^

Empirical experimental optimization of binding affinity can be laborious and costly, both in terms of time and resources.^18^ Molecular dynamics (MD) simulations, on the other hand, offer a valuable tool for the investigation of the interplay between stabilizing interactions, fluctuation correlations, and conformational variability at different levels of resolution and experimental conditions.^19–22^ *In silico* structural investigation of immunoglobulins and antigen-antibody complexes has been successfully employed with different objectives, which include the detailed description of the dynamics of CDR loops and the transitions between their conformational states,^23–25^ as well as the comprehension of structural rearrangements and allosteric modifications following antigen binding.^26–28^

Here, we employ atomistic MD simulations to investigate the internal dynamics of full-length pembrolizumab, a humanized IgG4 antibody used in immunotherapy, whose full structure has been experimentally solved29 (Figure 1). Pembrolizumab, whose commercial success rate is expected to profoundly impact pharmaceutical market in the next years, ^30^ is approved for the treatment of melanoma, lung cancer, head and neck cancer, Hodgkin’s lymphoma and stomach cancer.^31–33^ Its mechanism of action consists in binding to the programmed cell death protein 1 (PD-1), a 288-residues long receptor located on the membrane of T cells, B cells, and natural killer cells.^34^ PD-1 promotes apoptosis of the lymphocyte when activated by the programmed cell death receptor ligands PD-L1 and PD-L2, whose expression is upregulated in malignant cells.^35^ The large contact area between pembrolizumab and PD-1 hinders the binding of PD-L1 and PD-L2, thus preventing down-regulation of the anti-tumor activity of T cells.^36–38^

**Figure 1:**
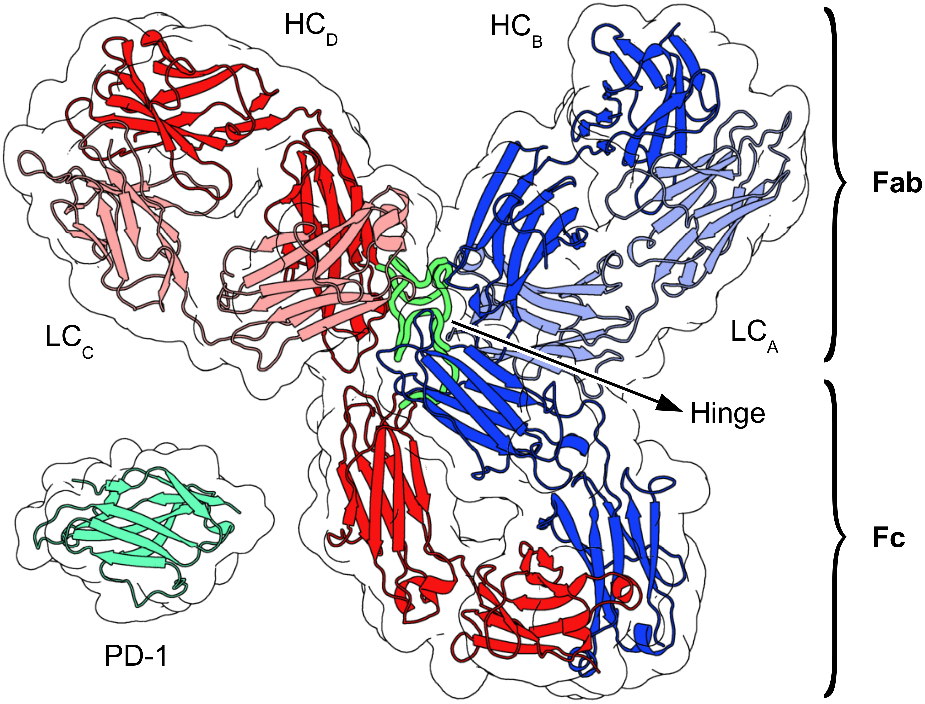
Graphical representation of the starting configuration of pembrolizumab employed in the MD simulations. LC stands for *light chain*, while HC stands for *heavy chain*. The antigen PD-1 is also represented.

We performed a total of 4 *μ*s of dynamics of the deglycosylated antibody, both in presence and in absence of the antigen PD-1. The analysis pipeline, which combines structural analysis, investigation of chemical interactions, and information theory-based measures of correlation, allowed us to rationalize at a residue-level the observed conformational dynamics. Our simulations highlighted the particular role of key residues of the hinge, resulting in an asymmetric behavior of the two hinge segments in pembrolizumab; moreover, an interplay between large-scale conformation and binding state is observed, and the residues allowing such information routing from the paratope throughout the antibody structure are identified. We believe that the residue-level elucidation of the complex dynamics of potent and highly selective mAbs – of which pembrolizumab is a notable example – in relation to their binding state may assist in the process of antibody engineering, for the rational design of novel, optimized therapeutic agents.

## Results and discussion

Pembrolizumab is a 1324-residues long therapeutic antibody, belonging to the IgG4 class. A schematic representation of it is given in Figure S3, where an index is associated to each structural domain; the corresponding residue ranges are listed in Table S2. As a first step, we focused on the analysis of conformations and the rationale behind domain motion in dependence of the binding state; to this aim, the results presented here are based on a comparison between the deglycosylated antibody alone (*apo* form) and bound to its antigen, the protein PD-1 (*holo* form). To allow for an unbiased comparison, the starting structure of the holo system was built on the same initial antibody conformation of the apo case; specifically, one Fab of pembrolizumab was replaced with the Fab-PD1 complex (PDB ID: 5GGS), after structural alignment on the antibody domain.

### The presence of the bound antigen restricts the range of conformations of the full-length pembrolizumab

To facilitate the analysis of antibody flexibility, the conformations sampled from MD simulations are collected in clusters on the basis of their structural similarity (Figure S4), as measured by the root-mean-square deviation (RMSD) matrix of C*_α_* atoms. For all systems, an RMSD threshold of 1.2nm is used. In the simulations of pembrolizumab alone, 6 conformational clusters are identified, while in the case of the holo system the same clustering protocol leads to the identification of only 4 clusters; in the presence of the antigen, a limited conformational variability is indeed observed, as apparent from the narrow range of radii of gyration spanned by these clusters when compared to the apo case (Figure 2). This is consistent with the visual inspection of the sampled conformations, whose representative structures are reported in Figure 2. A similar population shift towards a more uniform conformational distribution upon antigen binding was observed through MD simulations of an IgG1 antibody.^27^

**Figure 2:**
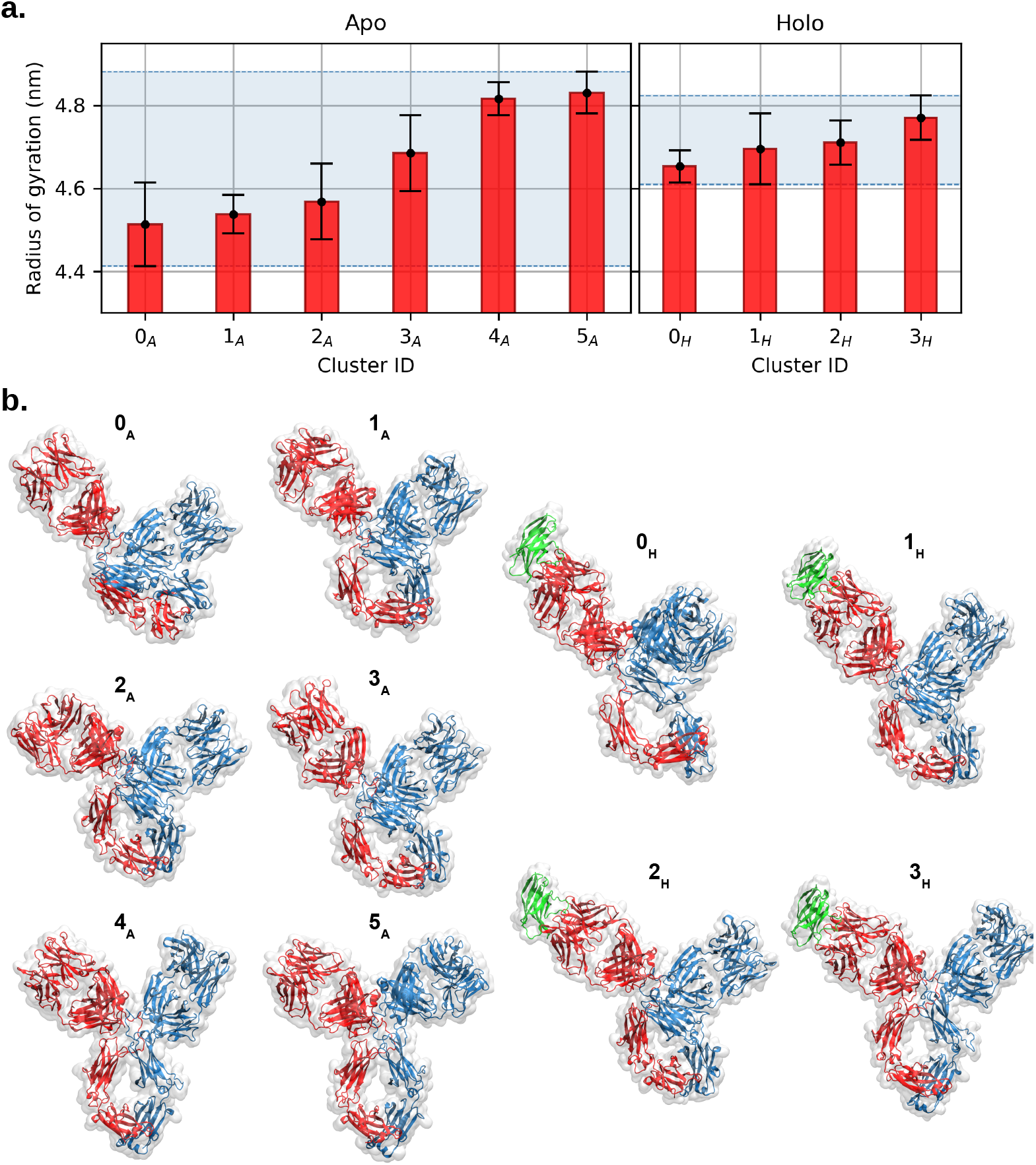
**a.** Mean and standard deviation of the antibody radii of gyration in the apo forms (*left*) and holo forms (*right*), averaged within each conformational cluster. In the holo case, the radius is computed for the antibody alone. The error bars correspond to the standard deviation within each cluster. The shaded areas correspond to the range of variability, which is greatly reduced in the holo case. **b.** Representative structures of pemrolizumab in the apo (A) and holo (H) forms, for each conformational cluster. Chains AB are in blue, chains CD in red, and the antigen PD-1 in green.

For each system (apo or holo), clusters are indexed according to increasing radii of gyration; in this way, clusters 0*_A_* and 0*_H_* are the ones grouping the most compact conformations of the apo and holo systems, respectively. At a large scale, conformational clusters differ mainly in the relative arrangement of the Fab and Fc domains, reflecting the extent of packing. Clusters 3*_A_* and 2*_H_* include the equilibrated experimental structures of the apo and holo systems, respectively. These clusters are intermediate, in terms of radius of gyration, between clusters 0*_A_*/1*_A_*/2*_A_* and 4*_A_*/5*_A_* for apo, and 0*_H_* / 1*_H_* and 3*_H_* for holo; this means that the simulations captured both tendencies of the antibody to shrink and to expand. Interestingly enough, the conformations that are not sampled by the holo system correspond to particularly low radii of gyration, namely those found in clusters 0*_A_*, 1*_A_*, and 2*_A_*. Compact conformations, particularly clusters 0*_A_*, 2*_A_* and 0*_H_*, present a higher structural stability, as measured in terms of RMSD with respect to the representative structure of the cluster (Figure S5).

The relative cluster population (quantified by the fraction of frames relative to the total number, Figure S6), shows a large unbalance between conformational states. The unbound antibody evolves towards a compact conformation for most of the simulation time, while the holo has a more symmetric distribution of configurations, associated to larger and smaller radii of gyration compared to the initial structure. Also in the holo case, however, the time spent in open conformations is a small fraction of the full simulation time. This result is in agreement with the experimental observation that the compact conformation is the most populated for IgG4 molecules in solution, as observed in ^15^N TROSY NMR and SAXS experiments.^29,39^ This conformational preference has been explained on the basis of the short hinge of IgG4 antibodies with respect to other IgG subclasses; IgG4 hinge has indeed a three amino acid deletion, compared to the one of IgG1.^40^ Another factor that can favour the compact arrangement is the tilted conformation of the CH2 domain in the Fc,^29^ which displays a rigid-body rotation of ≈ 120° relative to the position observed in the crystal structure of a truncated IgG4 Fc (PDB ID: 4C54). In the most compact conformations, the superposition of the pembrolizumab Fc and the one of 4C54 would lead to an overlap of CH2 and CL in Fab1 (Figure S7). The hypothesis that the CH2 domain rotation persists in solution was supported by a recent experimental observation of the cleavage of reduced pembrolizumab, mediated by the immunoglobulin-degrading enzyme from *Streptococcus pyogenes*, which binds the CH2-CH3 interface.^41^ The asymmetric orientation of the two CH2 domains can also explain the contact area between Fabs and Fc, reported in Figure S8. Here, the range of variability of the Fab1-Fc contact area greatly exceeds that of the Fab2-Fc pair, which remains low in the vast majority of the clusters; moreover, a significantly larger contact surface between Fab1 and Fc is observed in clusters 0*_A_* and 0*_H_*, with respect to the other cases. The rotated CH2 forms indeed a cavity that gives Fab1 a greater freedom to move, minimizing steric clashes with Fc. This is reflected also in clusters 4*_A_*, 5*_A_* and 3*_H_*, where the symmetric arrangement of Fabs with respect to Fc leads to a slightly larger Fab2-Fc contact area than Fab1-Fc; and in cluster 2*_A_*, 3*_A_* and 1*_H_*, where, although the Fc is clearly tilted toward the Fab1, the Fab1-Fc contact area is only slightly larger than the Fab2-Fc case. The large possibility of rearrangements of Fab1 with respect to the Fc is therefore one of the main determinants of the overall shape of pembrolizumab.

A comparison between the conformational states in apo/holo clusters is facilitated by the RMSD matrix in Figure S9. As expected, cluster 3*_H_* shares a low RMSD with the open conformations of clusters 4*_A_* and 5*_A_*, while clusters 1*_H_* and 2*_H_* are closer to the compact conformations in clusters 1*_A_*, 2*_A_*, 3*_A_*. The extremely compact conformation of cluster 0*_A_* and the peculiar cluster 0*_H_* are the ones sharing the lowest similarity with all the other clusters. 0*_H_* appears indeed as a singular conformation, where the major axis of Fab1 is perpendicular to the plane of Fab2 and Fc; once again, this movement is permitted by the rotated CH2 domain. Although such bent conformation of Fab1 is not directly observed in the apo case, visual inspection of the trajectory reveals that the variable region of Fab 1 in cluster 4*_A_* has the tendency to bend in a direction perpendicular to the plane of Fc-Fab2. This is confirmed by a principal components analysis (PCA) of the dynamics, performed on the C*_α_* atoms of the structures grouped in the conformational clusters (Section S2.1). The entity of the conformational change, however, is larger in 0*_H_* than in any apo cluster.

### Role of key residues in stabilizing the conformational states

The determinants of the observed conformations are identified, at a finer level of detail, through the calculation of correlations between residues, as quantified by the mutual information (MI) of C*_α_* atom fluctuations. MI, which captures both linear and non-linear contributions to amino acid displacements from a reference position, is used to build inter-residue networks of information pathways, which can be interpreted in the light of the inter-domain non-bonded interactions established in the course of the simulations. Table S3 and Table S4 list the residues involved in inter-domain contacts, including hydrogen bonds, salt bridges and hydrophobic interactions. These contacts are highlighted as important channels for information transfer by the results of the network analysis, and in particular by the calculation of edge betweenness (Figure S30 and Figure S31), which measures the centrality of a graph edge as the number of shortest paths crossing it.^42^

The most open conformations (clusters 4*_A_* and 5*_A_*) are stabilized by a persistent electrostatic interaction between side chains of ASP^788^ of Fab2 and LYS439 of Fab1, which results in a highly central edge of the interaction network; we can expect that mutation of one of these residues would destabilize the open state. Despite this common feature, stable contacts between Fab1 and Fc are absent in cluster 5*_A_*, while in 4*_A_* the two domains interact through electrostatic interactions. Moreover, the comparison of the non-bonded interactions between Fab1/Fab2 and Fc confirms the asymmetry of the contact distributions in the extended conformations: even though the molecule adopts an overall symmetric Y-shape, the larger number of interactions between Fc and Fab2 (with respect to Fab1) in clusters 4*_A_*, 5*_A_* and 3*_H_* supports the hypothesis that further approaching of the two domains would be impaired by steric clashes.

In the most compact conformations (clusters 0*_A_*, 1*_A_* and 0*_H_*), a key stabilizing role is played by ARG^435^ of Fab1, which emerges as a central network node and is involved in electrostatic interactions with several Fab2 residues (ASP^788^ and LYS^792^ in particular). The network representations of closed conformations reveal also the large number of edges crossing the Fab1-Fc contact surface in the compact arrangements. While the residues involved in these key interactions belong mainly to the CH2 domains of chain B, in the case of cluster 2*_A_* the surface of interaction is extended to distant regions of the Fc domain, namely the CH3 domain of chain B (through the backbone of MET^646^ and the side-chain of HIS^647^) and the CH2 domain of chain D (through the side-chains of THR^1215^ and LYS^1214^). The peculiar conformations in cluster 0*_H_* are stabilized by a number of high betweenness contacts taking place outside of the hinge; fundamental for the stability of this compact and highly interconnected state are the electrostatic interactions between side chains of LYS^856^-GLU^512^ in Fab2 and Fc, respectively, and between LYS^464^/ ARG^473^ in the AB loop of CH243 and Fab1 residues (specifically, hydrogen bonds LYS^464^-ASP^174^, ARG^473^-GLU^17^, and ARG^473^-PRO^15^.

These observations are strengthened by the analysis of communities in the MI-based network. The latter is divided into substructures (communities) with dense internal correlations, but sparse inter-community connections (see Section S1). Although the optimal community distribution closely reflects the natural subdivision of the antibody in structural domains, residues previously identified as promoting inter-domain connections fall within the same community (Figure S32 and Figure S33), thus supporting results from the investigations of non-bonded interactions and edge centrality. In addition, community analysis serves to detect, at a first level, connections between the hinge and the neighbouring domains; in this regard, a significant consistency is observed in the intermediate and fully open conformations of both apo and holo cases, where the hinge belongs to the same community as the CH1 domain of Fab2. Building on the high level of connection between the hinge and the rest of the molecule, a more detailed investigation of the role of the hinge in the overall confomational variability of the antibody is explored in the next section.

### Role of the hinge in the observed conformational variability

As noted above, the hinge displays significant correlations with other domains, both in the apo and holo systems, as apparent from the MI matrices (Figure S34 and Figure S35); such correlations appear particularly strong in pembrolizumab when compared to those reported in a previous study of an IgG1 antibody.^44^ In pembrolizumab, in fact, the hinge retains a complex network of interactions with nearby domains, thus functioning as more than just a flexible linker; a detailed account on the interactions involving the hinge residues, as emerging from network analysis and inspection of non-bonded contacts, is reported in Section S2.2. Given its role as interaction hub, in addition to allowing large-scale rearrangements, ^45,46^ the hinge region deserves therefore particular attention.

During our simulations, the hinge shows relatively small variations in the radius of gyration (Figure S36). Such result is expected on the basis of the presence of disulfide bonds between the two hinge segments, and of a number of transient intra- and inter-chain non-bonded interactions; among them, the most stable ones are the hydrogen bonds between the sidechain of SER^1100^ and the backbone of PRO^443^, and between the sidechain of TYR^1102^ and the backbone of PRO^446^ (Figure 3).

**Figure 3:**
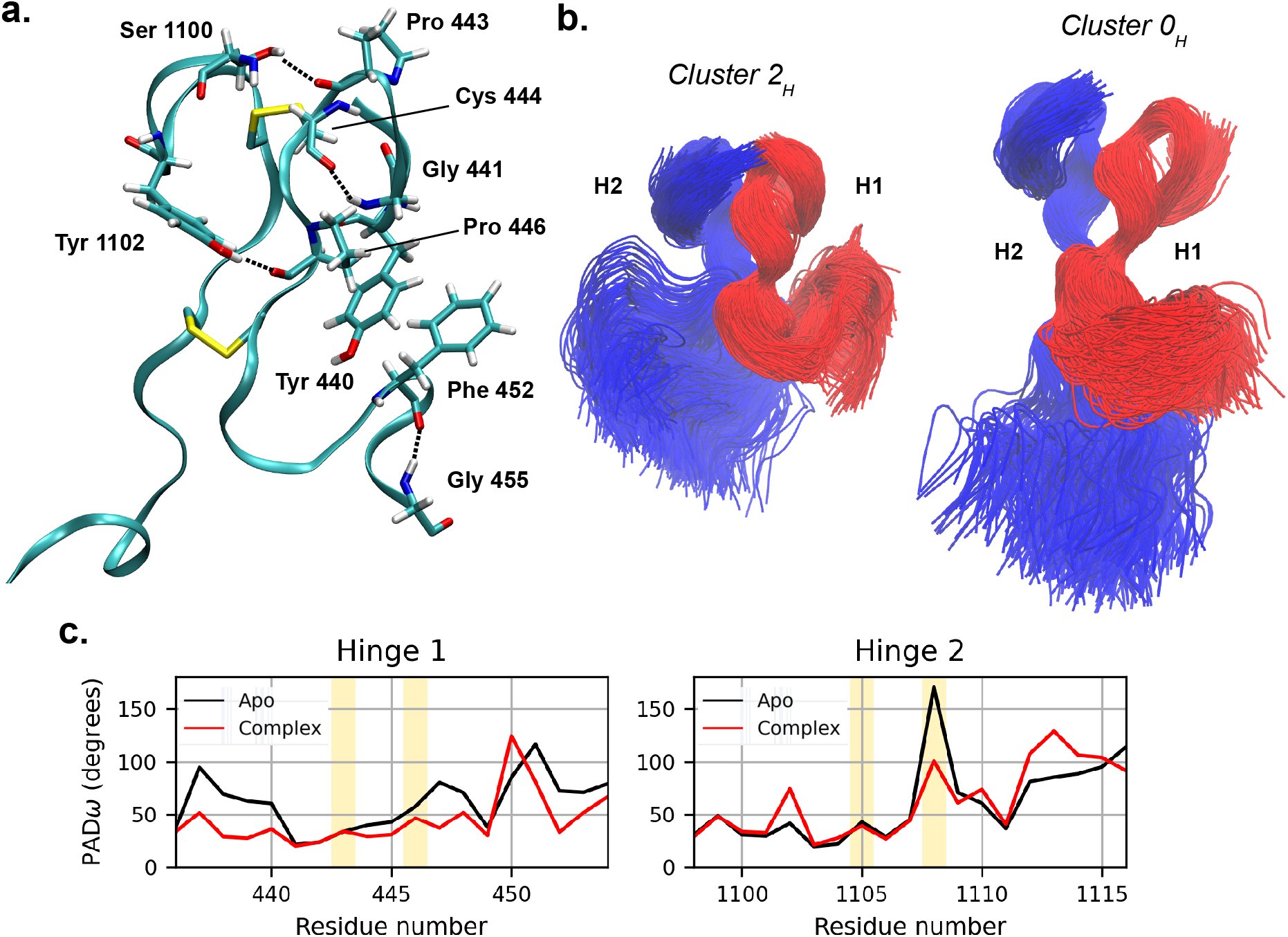
**a.** Interaction stabilizing the hinge conformation. The disulfide bonds between CYS^440^-CYS^1106^ and CYS^447^-CYS^1109^are also shown. **b.** Conformational ensembles of the hinge in the most compact (2*_H_*) and most extended (0*_H_*) clusters, after structural alignment on the cysteine residues. Chain B is represented in red, chain D in blue. The overall hinge shape is determined by the conformation of H2. **c.** Per-residue flexibility of hinge backbone, as quantified by PAD*_ω_* parameter. The yellow-shaded areas correspond to the cysteine residues forming inter-chain disulfide bonds. Particularly high is the backbone plasticity of the second cysteine in the hinge 2 of the apo antibody, which might result from the torsional stress imposed by the high conformational variability of the Fc relative to the Fab in the apo simulations.

The two hinge segments from chain B and chain D are highly asymmetric (Figure S37), with hinge 1 assuming significantly bent conformations (Figure 3). Although they are mostly attributable to the numerous non-bonded Fab1-Fc interactions, which prevent an extended conformation of chain B (see previous section), they are also stabilized by a number of non-bonded interactions, such as the stacking interaction between the side-chains of TYR^440^ and PHE^452^ and the hydrogen bonds between PHE^452^-GLY^455^ and GLY^441^-CYS^444^. Hinge 2 shows instead the largest degree of variability, and it appears to be the main determinant of the overall hinge shape.

Results from the previous sections highlighted a smaller range of conformations of the pembrolizumab molecule in presence of the antigen. In order to have a closer look at the residues responsible for the increased rigidity in the holo structures, we computed the PAD*_ω_* parameter (Figure 3), which measures the per-residue backbone plasticity through the variance of its torsional angles.^47^ PAD*_ω_* ranges from 0 to 180°; a higher value corresponds to a higher backbone flexibility. PAD*_ω_* values of hinge 2 (connecting Fab2, where the antigen is bound, and Fc) are very similar in the apo and holo cases, and for a few residues are even slightly higher in the bound state (the average 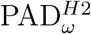 is 60° for apo and 62° for bound). However, hinge 1 (connecting Fab1 and Fc) is, on average, more rigid in the holo than in the apo form (the average 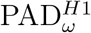 is 61° for apo and 44° for holo). Since the flexibility of hinge 1 is responsible for the relative movements of Fab1 and Fc, the calculated PAD*_ω_* values are in line with the restricted conformational variability observed in the bound conformations. RMSF computed on the hinge C^*α*^ atoms confirms the reduction of flexibility in hinge 1 with respect to hinge 2 when going from the apo to the bound state (Figure S38). Therefore, we suggest that the limited conformational variability of pembrolizumab in presence of the bound antigen is linked to a higher rigidity in the hinge region with respect to the apo case. Moreover, hinge 1 shares a higher MI with domains CH1 and CL of Fab2 than hinge 2 (Figure S34 and Figure S35), despite the latter being the direct extension of Fab2. Results from network analysis corroborate this observation: hinge 2 in the holo system does not include residues with high centrality, as opposed to the apo case (Table S5 and Table S6).

The hinge is not the only region of the holo antibody where a decrease in backbone plasticity is observed, with respect to the apo case; from the PAD*_ω_* analysis, this is true also for the binding site of Fab2 (as expected for the presence of the antigen) and residues 460-470 and 556 in the CH2 domain of chain B (Figure S39). Interestingly enough, this domain is indeed involved in high-betweenness contacts, as evidenced by the network analysis (Figure S30 and Figure S31), through highly central paths that put it in communication with Fab2. We notice here that a concurrent reduction in flexibility in the binding site, in hinge 1, and in the CH2 domain is not in contradiction with the observation that changes of conformational flexibility in antibodies follow Le Chatelier’s principle:^48,49^ upon binding, counteracting changes in rigidity and flexibility occur at distant sites. Figure S39 reveals indeed that a reduction in the value of PAD*_ω_* in the paratope of Fab2 is associated to an increased backbone plasticity in distant regions of the holo system, especially in the chain B of the Fc domain (residues 1132-1135, 1151, and 1203-1210).

### Binding modes of PD-1 elicit different degrees of correlation within the antibody

The results from the previous section suggest the presence of a communication channel between the antigen-bound Fab2 and hinge 1, leading to a transfer of information that might modulate an interplay between binding and conformational state. In this regard, different conformational clusters correspond to different binding site conformations, as quantified by the distributions of the RMSD between C*_α_* atoms in the paratope of the simulated system and the bound crystal structure^50^ (Figure S40). At a first level of distinction, the presence of the antigen largely reduces the displacements in the binding site, as expected; moreover, in the apo case, a correspondence is observed between the entity of the displacements and the compactness of the conformation, with more compact clusters shifted toward larger values of RMSD.

In order to better understand the interplay between the binding site and the rest of the pembrolizumab structure, we performed an investigation of the stabilizing interactions between the antibody and the antigen in each holo cluster, followed by a study of the correlations elicited by the ligand within the antibody.

The complex formed by Fab2 and the protein PD-1 is shown in Figure 4. The number of hydrogen bonds between the two molecules varies slightly among the distinct conformational clusters (Figure S41); large deviations are instead observed in the persistence of the salt bridge formed between side-chains of ARG^979^ of Fab2 and ASP^1377^ of PD-1 (Figure 4), which has been identified as a key contact for the stabilization of the complex.^50^ For all electrostatic interactions, the binding is strengthened when the antibody is in cluster 0*_H_*, resulting in large and stable values of the contact surface area between pembrolizumab and PD-1 (Figure S41). The RMSF of C*_α_* atoms in the antigen molecule, after structural alignment of the antibody variable region, reflects the strength of the interactions, with fluctuations in 0*_H_* that are approximately the half of those in the other clusters (Figure S42). Furthermore, the distribution of the antigen RMSD in each cluster, calculated with respect to the experimental structure, reveals a lower dispersion in cluster 0*_H_* (Figure S42). This particular stability might be correlated to the rigidity of the antibody structure in this conformation (Figure S5), and the corresponding absence of large-scale conformational changes perturbing the Fab/antigen complex. These observations are in agreement with the calculated values of binding enthalpies, obtained with the MM/PBSA approach^51^ and reported in Table S7. The values shown are relative to the simulation with the sole Fab in complex with the antigen, taken as the reference. In addition, for all of the conformational clusters, the simulation of the full-structure antibody results in a more stabilizing energy than in the simulation of the Fab alone, thus suggesting the importance of maintaining the full antibody structure in the studies of antibody/antigen binding free energies through MD simulation; a similar stabilizing effect was already observed upon inclusion in the simulation set-up of the constant region, with respect to the sole variable region.^52^

**Figure 4:**
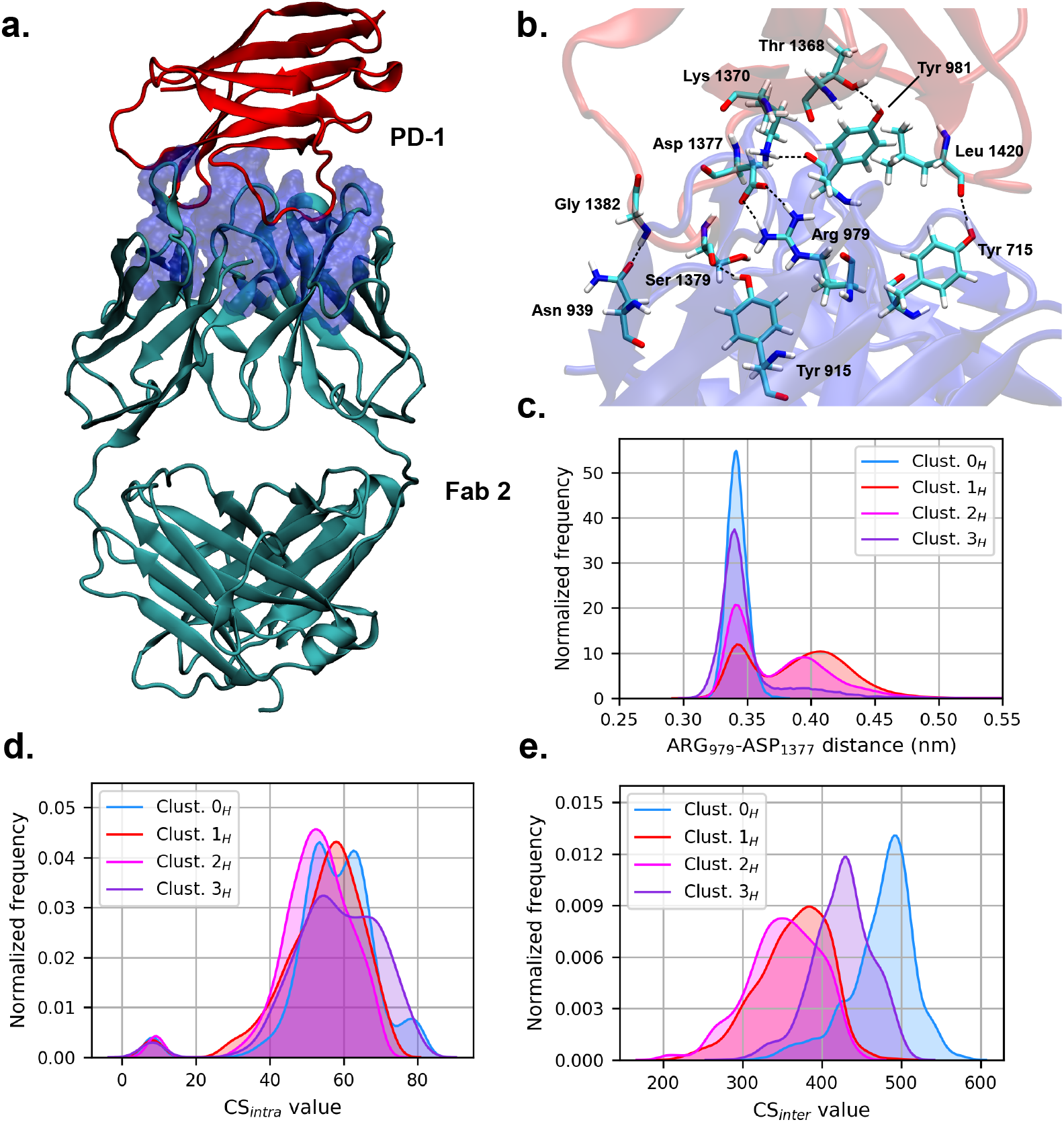
**a.** Pembrolizumab Fab2 (*cyan*) in complex with PD-1 protein (*red*). The paratope is represented as a blue surface. **b.** Residues involved in electrostatic interactions at the atigen/antibody interface. **c** Distance distributions between ARG 979 of Fab and ASP 1377 of PD-1. **d** Distributions of the intra- and inter- (**e**) domain correlation coefficients of antibody residues, for each conformational cluster of the holo state.

An ordering of conformational clusters similar to the one emerging from the strength of the binding interaction is reflected in the intensity of the residue-residue correlations within the antibody, as quantified by the generalized correlation coefficient (GCC);^53^ the latter corresponds to a normalized form of MI (Section S1), ranging from 0 (no correlation) to 1 (perfect correlation). Distributions of GCC values are shifted toward higher values in those clusters with the tightest binding (Figure S43), particularly in cluster 0*_H_*. For a more detailed inspection, we followed the example set in Palermo et al.,^54^ and we computed the correlation score (CS) for each residue *i* as the sum of GCC values with all the other antibody residues. This allowed us to distinguish *intra*-domain scores (Figure S44), where the summation extends to the residues belonging to the same structural domain, and *inter*-domain scores (Figure S45), which take into account only residues belonging to the other structural domains, excluding the one that includes residue *i*. From the distributions of intra- and inter-CS values shown in Figure 4, it is apparent that the change in correlations among the different clusters stems from an increased inter-correlation between the antibody domains in clusters 0*_H_* and, to a smaller extent, in cluster 3*_H_*. On the opposite, the intra-correlations do not show significant variations. In cluster 0*_H_*, in particular, the inter-correlations are significantly higher in Fab2 and in the chain B of Fc, with respect to the other clusters (Figure S45). This result can be explained on the basis of the tight binding between antigen and antibody in cluster 0*_H_*, and of its largely distributed communication network, as highlighted above.

Mutual information shows the highest correlation between the antigen molecule and all the antibody domains in cluster 0*_H_* (Figure S46), especially in Fab1. To further investigate the interplay between paratope and such peculiar antibody conformation, simulations of pembrolizumab in the apo form were started from the representative structure of cluster 0*_H_*, after removal of PD-1. Three 100ns-long replicas were performed to allow residues in the paratope to relax to new equilibrium conformations, while the antibody retains an overall conformation closely similar to the starting one. As shown in Figure S47, the RMSD distribution of the residues in the binding site overlaps with the one obtained from the apo cluster with the highest structural similarities, namely cluster 1*_A_* (Figure S9); in addition to a large network of interactions, as observed in the case of the other compact apo clusters, the latter is characterized by particularly strong contacts between the Fab domains (Table S3). A similar transition of the binding site conformation is not obtained in the case of the other holo clusters, thus suggesting once again a complex interplay between the large-scale conformation of the antibody and the antigen binding site, as elicited by the highly extended correlation network of pembrolizumab in conformational states compatible with cluster 0*_H_*.

## Conclusions

While impressive progress has been made in the experimental characterisation and manipulation of antibodies, a detailed, atomistic investigation of their properties is still incomplete. This is particularly true for the interplay between the molecule structure and its dynamics, which is extremely rich and varied, as several studies have recently shown.^27,55–57^

If, on the one hand, the specific sequence of the CDR plays the most prominent role in the selectivity and binding affinity of the antibody, on the other hand the observation that a modulation of the binding strength can be effected though mutations in distal sites shows that the internal mechanics of these molecules can be extremely complex.^16,17^ This is particularly true for IgG4 antibodies such as pembrolizumab, whose short hinge reduces the degree of flexibility and tightens the interactions among the various domains.

In this work, we have made use of atomistic MD simulations and information-theoretical analysis methods to elucidate the relation between the large-scale arrangement of pembrolizumab and the stability of the antigen binding. The analysis pipeline employed has allowed us to highlight a substantial conformational variability, a quality rather different from that one might observe in the case of loosely-connected rigid bodies. On the contrary, a complex pattern of structural arrangements, intramolecular communication pathways, and binding strength has emerged.

In accordance with experimental results, we observed that the antibody is prevalently found in a compact, asymmetric shape. This particular arrangement can be explained on the basis of the short hinge, which limits the conformational space accessible to Fab arms, as well as the rotated CH2 domain of chain B.

Furthermore, the spectrum of structures sampled by the molecule is modulated by the binding state: the large conformational variability of the apo case is substantially restricted when the antibody is complexed with the antigen. Similarly, the flexibility of the hinge (hinge 1 in particular) is reduced in this latter case, with a high degree of correlation emerging between the binding site on Fab2 and hinge 1.

The binding site was shown to bear strict ties with the rest of the molecule in general. The analysis of the distributions of the apo binding site RMSD, computed taking the experimentally resolved holo structure as a reference, highlighted a correlation between the binding site and the large-scale arrangement of the molecule as a whole.

The binding of the PD1 antigen to Fab2 is, in general rather stable in all holo state simulations; however, the strength of the binding is different across the various conformational clusters. The trend follows the intensity of the inter-domain correlations not only within the antibody, but also between the antigen and the antibody itself; a particularly strong affinity is observed in a specific configuration in which Fab1 is bent toward Fab2 and Fc. The tight binding and high correlations in this cluster suggest an interplay between the binding mode and the intensity of correlations, which manifest themselves in an extended network of interactions.

The results reported in this work return a picture of antibodies as extremely complex molecules, with a rich pattern of structural and dynamical features. The analysis protocol here applied to pembrolizumab is completely general, which enables its widespread application to other antibodies, with the objective of acquiring an ever deeper understanding of their inner life and, in turn, providing sharper tools for the manipulation of these molecules for medical and technological purposes.

## Methods

The crystallographic structure of pembrolizumab in the apo form (Ab1, PDB ID: 5DK3)^29^ was used as a starting structure for the MD simulations, after modelling of the missing residues (Section S1). The system in the holo form (Ab2) was obtained by replacing one Fab in Ab1 with the crystallized structure of Fab-PD1 complex (PDB ID: 5GGS),^50^ after structural alignment. The Fab-fragment alone bound to the PD-1 (Ab3) was also simulated, in order to assess possible differences in the interaction with the antigen with respect to the full-length antibody.

MD simulations were performed with the Gromacs 2018 software.^58,59^ The amber99SB-ILDN force field^60^ was used to define the topology of the systems. The protein was solvated in a box of TIP3P water molecules^61^ and the charge was neutralized with Cl^−^ and Na^+^ ions at physiological concentration (150 mM). The final number of atoms was 571932 in Ab1, 618265 in Ab2, and 193009 in Ab3; of these, the number of protein atoms was 20218, 21976 and 8317, respectively. Energy minimization was performed until the maximum force reached the value of 500 kJ mol^−1^ nm^−1^. NVT and NPT equilibrations were performed for 100 ps each using the Velocity-rescale thermostat^62^ and the Parrinello-Rahman barostat.^63^ Temperature was set at 300 K and pressure at 1 bar, and the time constants were set to *τ*_*t*_ = 0.1 ps and *τ*_*p*_ = 2 ps. Constraints were applied to hydrogen-containing bonds through the LINCS algorithm.^64^ A cutoff of 10 Å was used for Van der Waals and short-range Coulomb interactions; long range electrostatics was treated with Particle Mesh Ewald. Ab1 and Ab2 were simulated for a total of 2 *μ*s each, in 4 independent replicas of 500 ns. Ab3 was simulated for 500 ns. Additional simulations of the apo state were started from the representative holo conformations, after removal of the antigen; the structures were solvated and equilibrated using the above-mentioned procedure, and three 100ns-long replicas were simulated for each system.

Clustering of the simulation frames according to structural similarity was performed with an in-house script using hierarchical clustering (UPGMA).^65^ Calculation of atomic displacement, residue fluctuations, interatomic distances and contact surface areas, as well as principal component analysis, were performed using Gromacs 2018 utilities, while calculation of MM/PBSA energy was performed with the software *g_mmpbsa*.^51^ The number of hydrogen bonds was computed with VMD,^66^ setting an angle cutoff of 25° and a maximum donor-acceptor distance of 3Å. Mutual information, generalized correlation coefficient and correlation score were computed with in-house scripts, and network analysis was performed using the scripts produced by Melo *et al.*^67^ For a detailed description of the methods used for data analysis, the reader is referred to the Supplementary Information.

## Data availability

The raw data produced and analysed in this work are freely available on the Zenodo repository https://doi.org/10.5281/zenodo.5018432.

## Acknowledgement

The authors thank Emiliano Biasini and Attilio Vittorio Vargiu for an insightful reading of the manuscript. This project received funding from the European Research Council (ERC) under the European Union’s Horizon 2020 research and innovation program (Grant 758588).

## Author contributions

RP designed the project; TT and MR performed the simulations and the analyses; all authors contributed to the interpretation of the results and to the writing of the manuscript.

## Notes

The authors declare no competing financial interest.

## Supporting Information Available

The Supporting Information is available free of charge.

- Additional methods; additional data and figures, including: sequence intervals for each structural domain of pembrolizumab; schematic representation of pembrolizumab; clustering of simulation frames; percentage of simulation time in conformational clusters; Fab/Fc contact area; RMSD distributions of full antibody; RMSD between clusters; comparison between pembrolizumab and 4C54 structures; principal component analysis for each cluster; inter-domain contacts; graphical representation of the networks; community network repartitions; mutual information averaged within structural domains; description of contacts in the hinge region; radius of gyration of the hinge; solvent-accessible surface area of hinge; PAD*_ω_* parameter; RMSF of hinge; RMSD of binding site; number of hydrogen-bonds; antigen-pembrolizumab contact area; RMSF and RMSD of antigen; MM/PBSA calculations of antigen/pembrolizumab binding; distributions of the generalized correlation coefficient; intra- and inter-domain correlation score; RMSD values of binding site in clusters 0*_H_*, 1*_A_*, and 0*_H_* after antigen removal; antigen-pembrolizumab mutual information.

