## Supplemental material for "How Communication Pathways Bridge Local and Global Conformations in an IgG4 Antibody: a Molecular Dynamics Study"

* Contributed equally to this work

### S1 Methods

**System preparation** The crystal structure of the full-lenght pembrolizumab (PDB ID: 5DK3) [1] presented missing residues in the hinge region (residues 448-450 and 1110-1115), which were modelled with Chimera [2] using an IgG2a antibody (PDB ID: 1IGT [3]) as a template. Glycans were removed from the crystal structure, so as to isolate the role of structural features of the antibody itself in the emerging dynamics. The bound system was obtained by replacing one Fab in the apo system with the crystallized structure of Fab-PD1 complex (PDB ID: 5GGS [4]), after structural alignment of the antibody domain. The same structure was used for the simulation of the Fab-fragment alone bound to the PD-1. MD simulations were performed as described in the Main text.

**Principal components analysis** Principal component analysis (PCA) was performed for each cluster using $C_\alpha$ atoms positions. Calculation, diagonalization and analysis of the covariance matrices were performed using Gromacs tools *gmx covar* and *gmx anaeig* [5]. In order to visualize the direction of movements captured by the eigenvectors, porcupine plots were generated using the extreme projections on the first three principal component, and visualized with VMD. The direction of the arrow in each $C^\alpha$ atom represents the direction of motion, while the length of the arrow characterizes the movement strength.
In order to tell whether two structural domains $\mathcal{A}$ and $\mathcal{B}$ move in the same direction in a cluster, we computed the following quantity for each of the first three principal components:

$$q_k^{AB} = \frac{1}{NM} \sum_{i \in \mathcal{A}} \sum_{j \in \mathcal{B}} \mathbf{v}_i \cdot \mathbf{u}_j \tag{1}$$

Here, $\mathbf{v}_i$ and $\mathbf{u}_j$ are vectors of the $k$-th mode, relative to the residues $i$ and $j$, and $N$ and $M$ are the numbers of residues in domains $\mathcal{A}$ and $\mathcal{B}$, respectively. Results for each mode are combined, and weighted by their corresponding eigenvalues $\lambda_k$:

$$Q^{AB} = \frac{\sum_{k=1}^{3} q_k^{AB} \lambda_k}{\sum_{k=1}^{3} \lambda_k} \tag{2}$$

Each entry $Q^{AB}$ in the resulting matrix refers to a couple of structural domains. Positive values correspond to parallel movements, while negative values to motions in an antiparallel direction.

**MM/PBSA calculations** In the MM/PBSA approach [6], the binding free energy $\Delta G_{bind}$ between the antibody (Ab) and the antigen (Ag), namely

$$\Delta G_{bind} = G_{complex} - G_{Ab} - G_{Ag} \tag{3}$$

is written as a sum of different contribution:

$$\Delta G_{bind} = \Delta H - T\Delta S = \Delta E_{MM} + \Delta G_{sol} - T\Delta S \tag{4}$$

Here, $\Delta E_{MM}$, $\Delta G_{sol}$ and $\Delta S$ are the changes in the gas-phase molecular mechanics energy, solvation free energy, and conformational entropy upon binding, respectively. More specifically,

$$\Delta E_{MM} = \Delta E_{int} + \Delta E_{ele} + \Delta E_{vdW} \tag{5}$$

$$\Delta G_{sol} = \Delta G_{PB} + \Delta G_{SA} \tag{6}$$

$\Delta E_{MM}$ includes the changes in the internal energy $\Delta E_{int}$ (due to bonded interactions), electrostatic energies $\Delta E_{ele}$, and the van der Waals energies $\Delta E_{vdW}$. $\Delta G_{sol}$ is the sum of the electrostatic solvation energy $\Delta G_{PB}$ and the nonpolar term $\Delta G_{SA}$ between the solute and the continuum solvent. $\Delta G_{sol}$ is calculated using the Poisson-Boltzmann model, while the nonpolar energy is estimated on the basis of the accessible surface area of the proteins.

In our calculations, we neglected the changes in conformational entropy, focusing therefore on the enthalpic contributions to the binding energy. These were estimated with the *g_mmpbsa* tool [7]. A convergence analysis was performed to estimate the appropriate number of frames (Figure S1), which are extracted from the trajectory at regular intervals. The average values and the corresponding errors are obtained with a bootstrap analysis. With ∼1500 frames, the result is fully converged with an acceptable uncertainty.

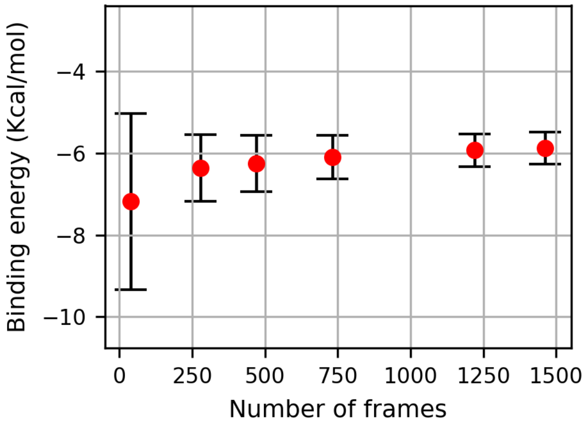


Figure S1: Convergence of the binding energy computed with the MMPBSA method as a function of the number of frames employed for the calculations.

**Correlations** Mutual information (MI) was calculated with as a measure of inter-residue communications. MI is defined as following:

$$MI_{ij} = \iint dx_i dx_j \; p(x_i, x_j) \log \frac{p(x_i, x_j)}{p(x_i)p(x_j)} \tag{7}$$

where $x_i$ and $x_j$ are the displacements of the atoms with respect to their average positions, $p(x_i)$ and $p(x_j)$ are the probability functions of finding the $i$-th or $j$-th atoms with a displacement equal to $x_i$ or $x_j$, $p(x_i, x_j)$ is the joint probability function. The calculation was performed with in-house scripts, and the displacement was divided in 100 discrete bins. From the mutual information, the generalized correlation coefficient ($GCC$) is computed as:

$$GCC_{ij} = \sqrt{1 - \mathrm{e}^{-2MI_{ij}/3}} \tag{8}$$

$GCC$ represents a measure of normalized MI, ranging from 0 (no correlation) to 1 (perfect correlation) [8]. Calculations were performed through in-house scripts.

GCC was used to calculate the correlation score (CS). For each residue $i$, $CS_i$ is computed as the sum of the generalized correlation coefficient values between residue $i$ and the other protein residues:

$$CS_i = \sum_{j \neq i} GCC_{ij} \tag{9}$$

In the case of the *intra*-domain $CS$, the sum extends to the residues belonging to the same structural domain as residue $i$; in the case of the *inter*-domain $CS$, the summation takes into account only residues belonging to all the other structural domains, excluding that of residue $i$.

**Network analysis** Protein networks were defined as sets of interconnected nodes centered on the $C^\alpha$ atoms. The total number of nodes of each system corresponds therefore to the number of residues. A couple of nodes is considered connected by an edge if any heavy atoms of the two residues is within a distance of 4.5 Å for at least 75% of the simulation time. These cutoffs were selected after a convergence study based on the Community Repartition Difference (see below).
Each edge is weighted according to the generalized correlation coefficient measure; specifically, the weight of the edge between nodes $i$ and $j$ is defined as:

$$w_{ij} = -\log\left[GCC_{ij}\right] \tag{10}$$

where $GCC_{ij}$ is the generalized correlation coefficient (equation 8).
The network analysis was performed with the Python implementation found in Melo et al. [9], which makes use of the NetworkX package [10] and is optimized using Cython[11] and Numba [12]. In the weighted networks, the sets of communities were identified using the Girvan-Newman algorithm [13, 14]. As a measure of the quality of the community structure, the modularity parameters $Q$ was calculated (Table S1)). $Q$ represents the difference in probability of intra- and inter- community connections for a given community repartition, and is defined as:

$$Q = \sum_i \left(e_{ij} - a_i^2\right) \tag{11}$$

where $e_{ij}$ is the fraction of edges that links nodes in community $i$ to nodes in community $j$, and $a_i = \sum_j e_{ij}$ is the fraction of edges from all communities that connect to nodes belonging to community $i$. The range of modularity values is between 0 and 1. Values close to 1 identify high-quality community structures, favouring connections intra-community with respect to inter-communities ones.

Table S1: Modularity of the community structures

| System | Cluster ID | Modularity |
|---|---|---|
| Apo | 0 | 0.8506 |
| | 1 | 0.8486 |
| | 2 | 0.8525 |
| | 3 | 0.8475 |
| | 4 | 0.8474 |
| | 5 | 0.8548 |
| Complex | 0 | 0.8517 |
| | 1 | 0.8557 |
| | 2 | 0.8575 |
| | 3 | 0.8585 |

All clusters from apo and complex simulations show similar values of modularity, which is very high in all the cases (Table S1). The above-average values (which are usually found between 0.4-0.7 for protein networks) can be explained on the basis of the natural partition of antibodies in structural domains.

Moreover, communities were used as a mean to evaluate the suitability of the cutoffs defining the edges. In order to evaluate the effect of the choice of the cutoffs on the resulting communities, the Community Repartition Difference (CRD) is calculated. The CRD between two network repartitions $c_1$ and $c_2$ is defined as:

$$CRD(c_1, c_2) = 1 - \frac{\sum_{n_i,n_j} z(n_i, n_j, c_1) z(n_i, n_j, c_2)}{\sum_{n_i,n_j} z(n_i, n_j, c_1)} \quad (12)$$

where $z(n_i, n_j, c_k)$ is defined as 1 if nodes $n_i$ and $n_j$ belong to the same community in a given network partition $c_k$, and 0 otherwise. A value of CRD equal to 0 indicates identical repartitions, while a value equal to 1 corresponds to the case of totally different communities. We computed CRD values for several combinations of distance/frame cutoffs, with respect to the reference case of 4.5 Å distance cutoff and 75% frame cutoff (Figure S2). These two values, which are in the range commonly used for protein network analyses [9], are the ones chosen for this study. In all cases, the value of CRD is below 0.2, indicating that a change of the parameters within the range investigated does not lead to substantial changes in the resulting community repartitions.

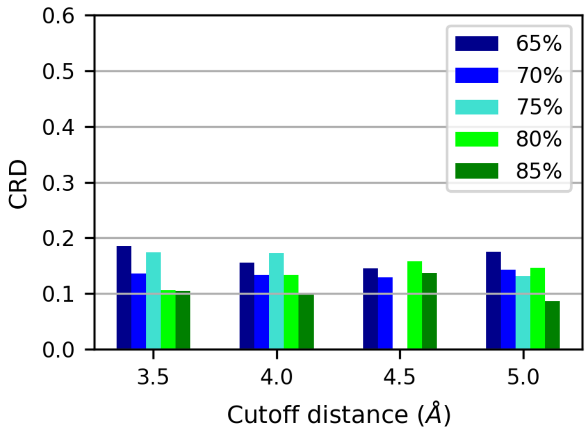


Figure S2: Community Repartition Difference (CRD) computed for different choices of the parameters defining the edges of the network. The "distance cutoff" (ranging here from 3.5 Å to 5.0 Å defines the maximum distance for which two atoms are considered in contact, while the "frame cutoff" (ranging here from 65% to 85%) defines the percentage of frames in which the contact is formed. The values reported in the plot refer to the CRD between each of the possible combination of the two cutoffs and the reference case with 4.5 Å distance cutoff and 75% frame cutoff.

Finally, edge betweenness was used to measure the importance of the edge for communication within the network. Edge betweenness, which is defined as the number of shortest pathways that cross the edge, was calculated with the commonly used Floyd-Warshall algorithm [15, 16].

**Images** Images of the proteins were produced by using VMD [17] and Protein Imager [18], and the graphs were made with python libraries.

### S2 Additional results

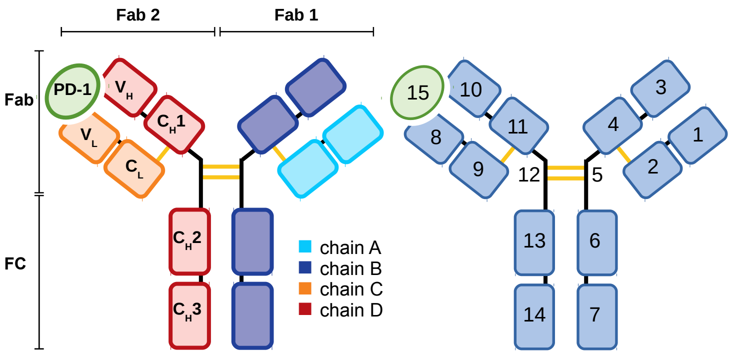


Figure S3: Schematic representation of the antibody pembrolizumab in complex with antigen PD-1, with the indication of the name of each structural domain. Chains C and D correspond to chains F and G in the original PDB file, respectively [1]. The numbering on the right corresponds to the indices used in the correlation matrices.

Table S2: Subdivision of the pembrolizumab/PD-1 complex into structural domains.

| Fragment name | Domain ID | Residues range |
|---|---|---|
| Fab 1 | 1 | 1-112 |
| | 2 | 113-218 |
| | 3 | 219-340 |
| | 4 | 341-438 |
| Hinge 1 | 5 | 439-456 |
| Fc 1 | 6 | 457-558 |
| | 7 | 559-662 |
| Fab 2 | 8 | 663-774 |
| | 9 | 775-880 |
| | 10 | 881-1001 |
| | 11 | 1002-1099 |
| Hinge 2 | 12 | 1100-1117 |
| Fc 2 | 13 | 1118-1225 |
| | 14 | 1226-1324 |
| PD-1 | 15 | 1325-1438 |

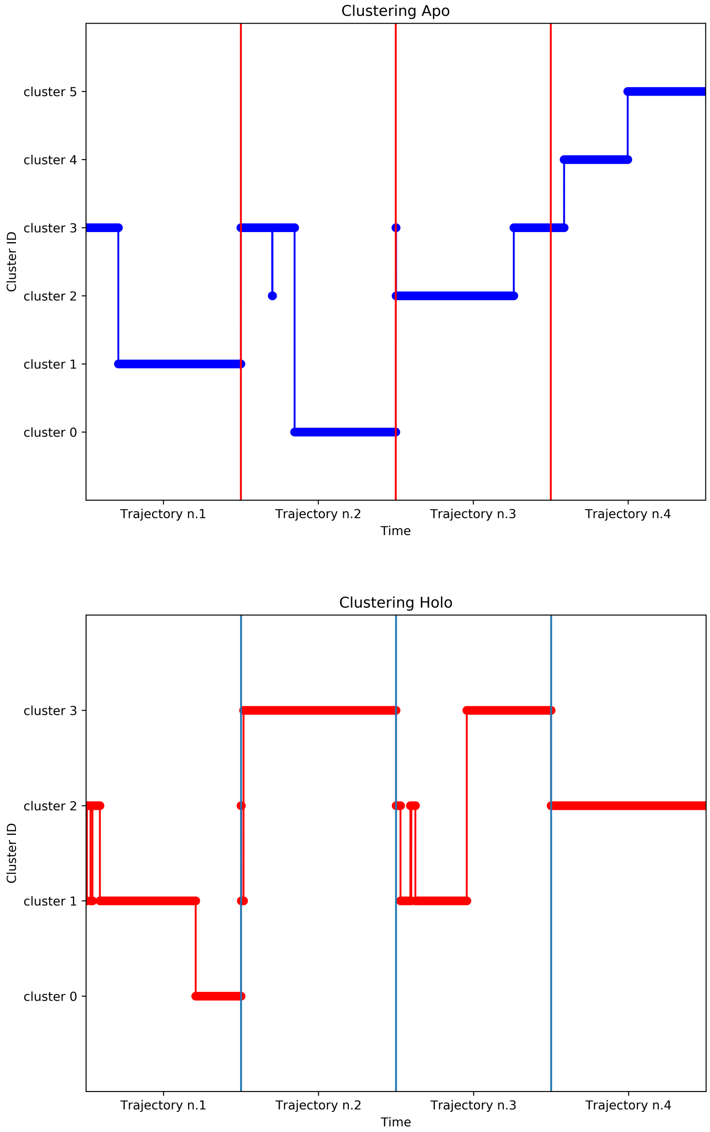


Figure S4: Timeline of the clustering assignment, in the apo (*top*) and holo (*bottom*) systems.

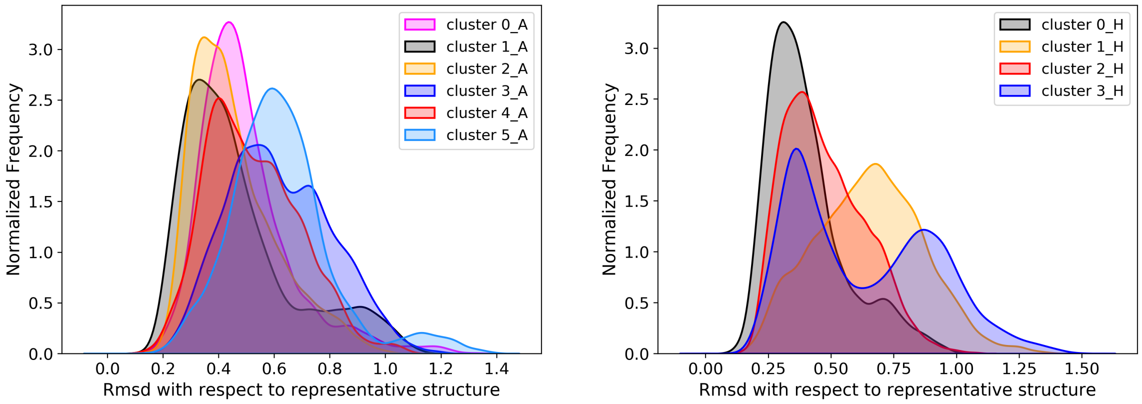


Figure S5: Rmsd distribution of antibody structure, for each cluster with respect to the representative conformation. Left: Apo case. Right: Holo case.

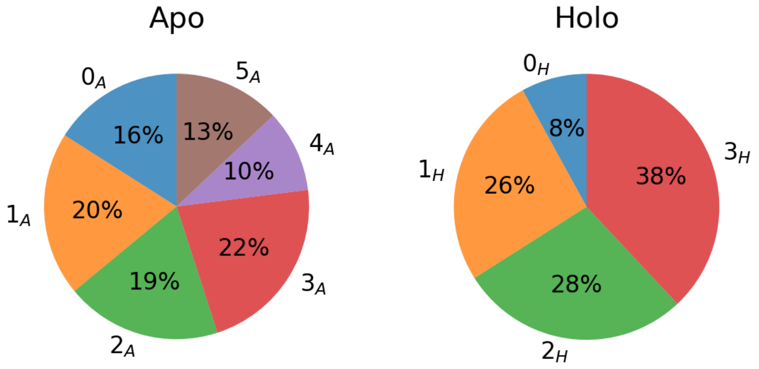


Figure S6: Overall occurrence of the conformational clusters in the apo and holo states.

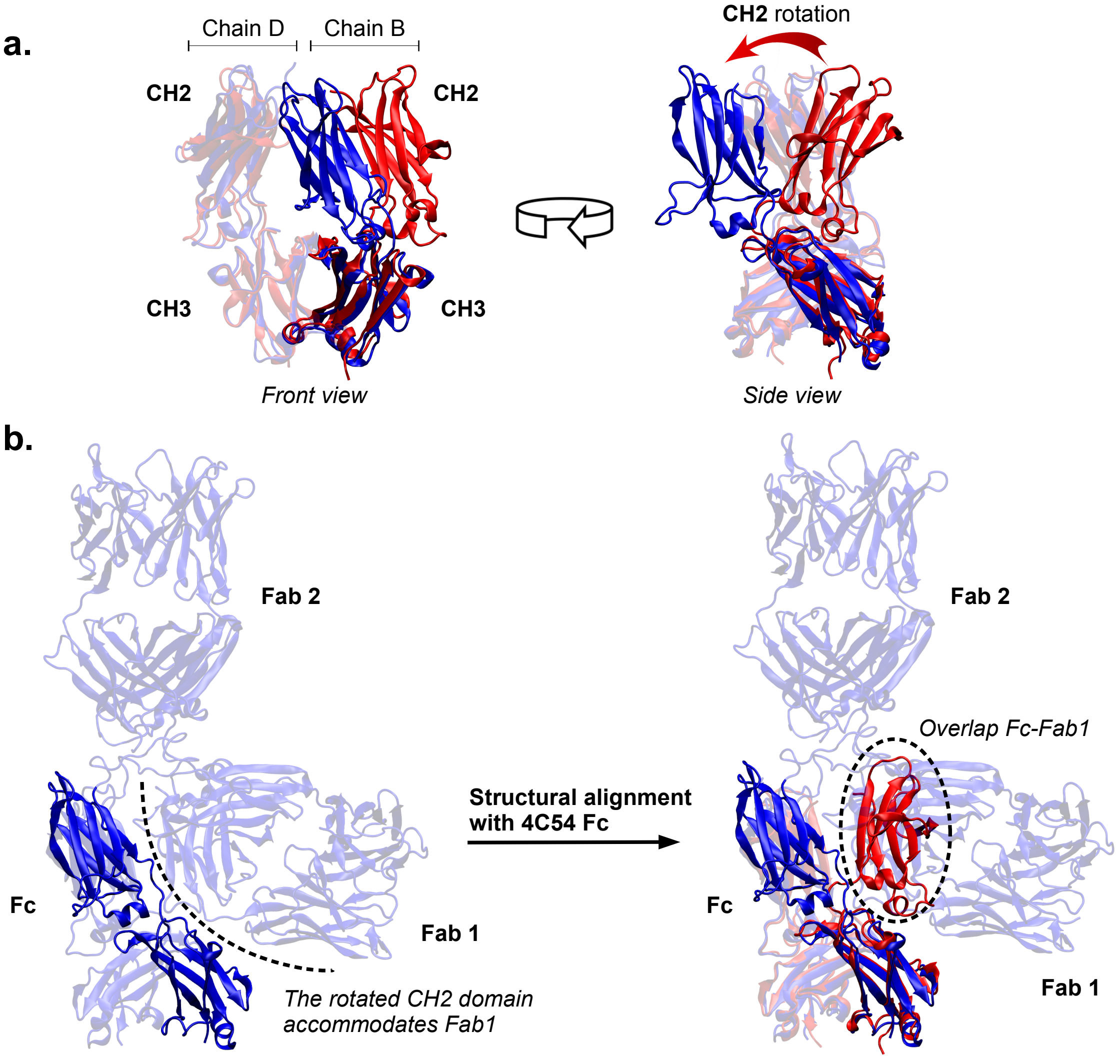


Figure S7: **a.** Comparison of Fc domains from pembrolizumab in the compact conformation (representative structure of cluster $0_A$, *blue*) and from an isolated IgG4 Fc (PDB ID: 4C54, *red*), after structural alignment. Chain B is solid color, while chain D is semi-transparent. **b.** Role of the rotated CH2 domain of pembrolizumab in the compact conformations. The close contact between Fab1 and Fc observed in the simulations would not be possible without the peculiar CH2 position; the conformation of CH2 found in 4C54 would lead to an overlap between Fc and Fab1.

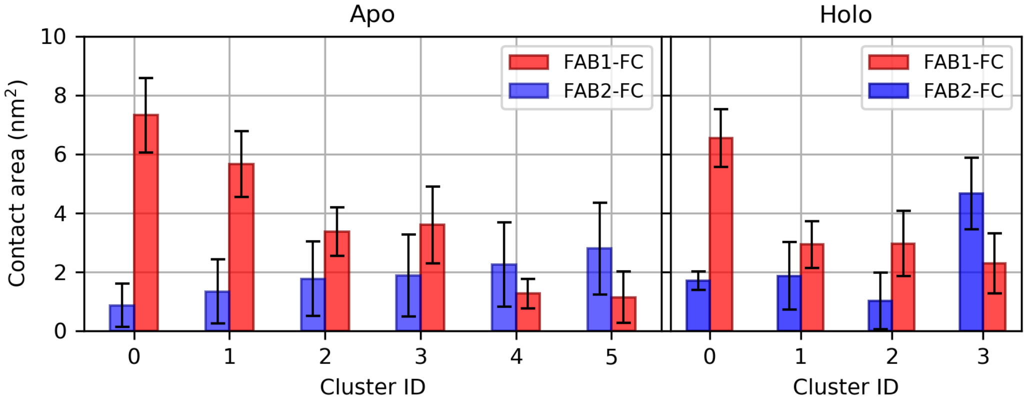


Figure S8: Average contact area between the Fab and the Fc domains, for each conformational cluster in the apo (*left*) and holo (*right*) states.

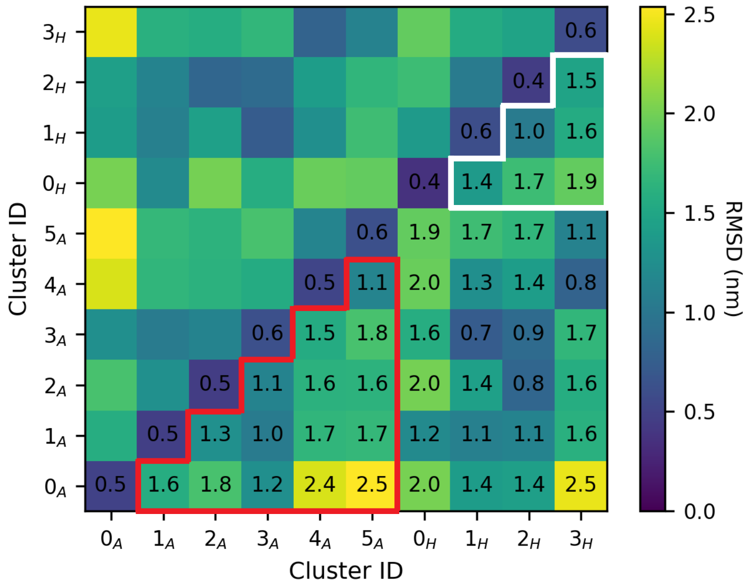


Figure S9: Average RMSD between frames belonging to each pair of clusters. The RMSD has been computed on all the $C^{\alpha}$ atoms of the antibody after structural alignment. Cells within the red border compare clusters of the apo system, while cells within the white border refer to the holo system.

### S2.1 Principal components analysis

#### Cluster $0_A$

Fab1 and Fc move apart from Fab2, while Fab1 and Fc remain in close contact during the movement, giving rise to a large contact surface. Fc is rotating in such a way that CH2 of chain B is moving apart from CL of Fab2.

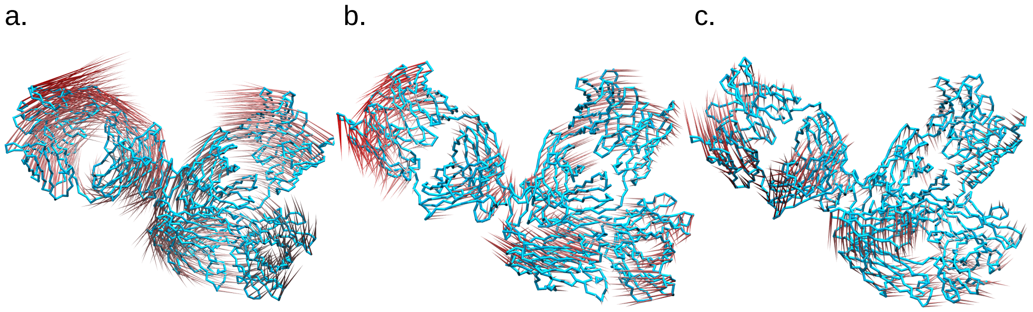


Figure S10: Porcupine representations of modes 1 (**a**), 2 (**b**) and 3 (**c**).

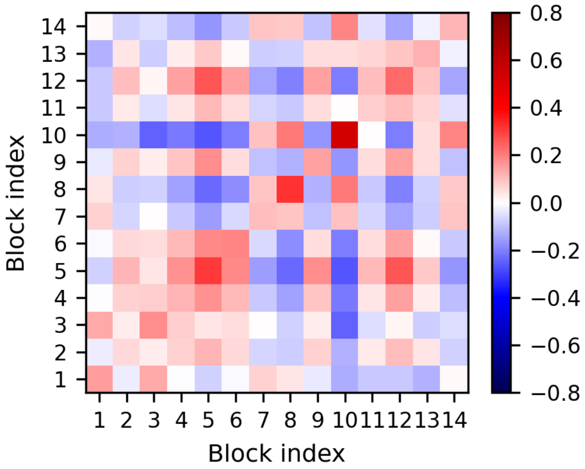


Figure S11: Collinearity of the structural domains in the first three modes.

#### Cluster $1_A$

We observed a high mobility of the variable region of Fab2, in particular VH. The dynamics is similar to the one of cluster $2_A$, but this time the CH3 domains tend to move in the same direction of the variable domains of Fab2.

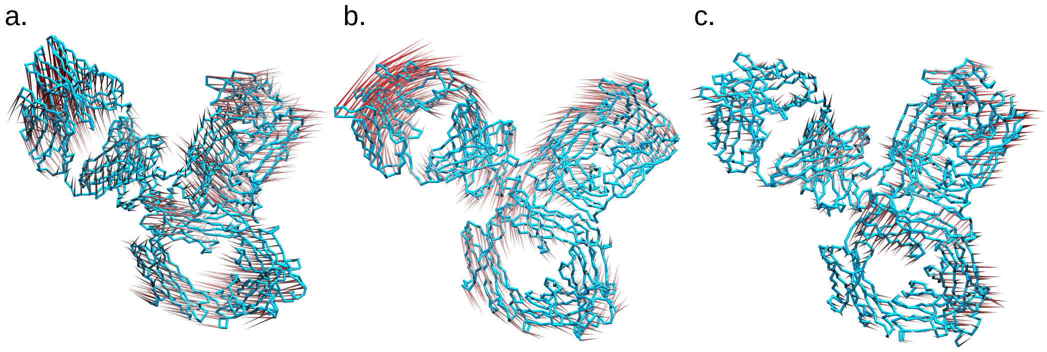


Figure S12: Porcupine representations of modes 1 (**a**), 2 (**b**) and 3 (**c**).

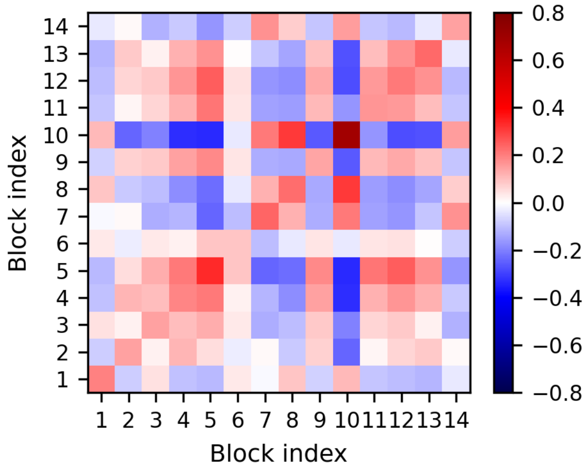


Figure S13: Collinearity of the structural domains in the first three modes.

### Cluster $2_A$

A deformation of Fab2 takes place: the variable domains of Fab2 move in a strongly anticorrelated motion with respect to the rest of chain D (whose directions of motion are highly parallel), and to the hinge of chain B.

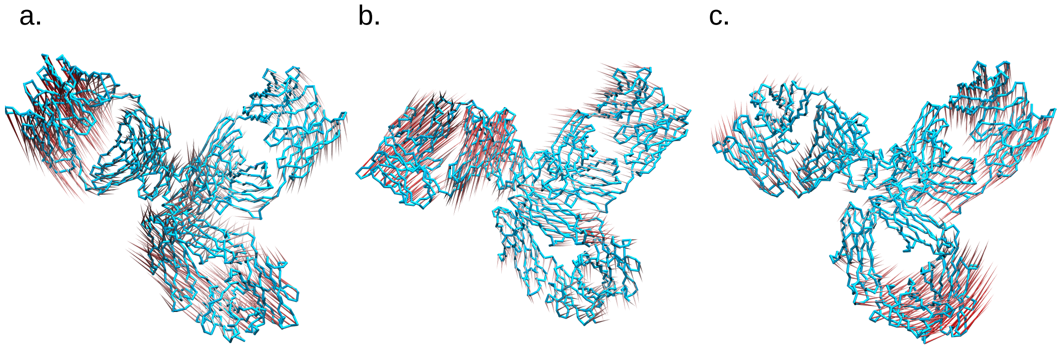


Figure S14: Porcupine representations of modes 1 (**a**), 2 (**b**) and 3 (**c**).

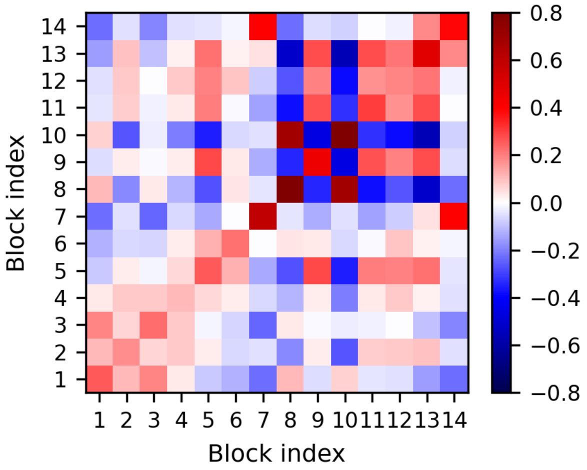


Figure S15: Collinearity of the structural domains in the first three modes.

#### Cluster $3_A$

Rotation of Fab2 takes place until the CL is in contact with the Fc (indeed, this is the only one among the compact clusters in which atomistic non-bonded interactions are formed between residues belonging to these two domains). This happens by means of parallel motions of the CH3 domains and the variable domains in Fab2. At the same time, the distance between the lower hinge segments is reduced, inducing a rotation of Fab1 in the opposite direction of Fab2. This movement of Fab1 is parallel to that of the remaining domains in chain D.

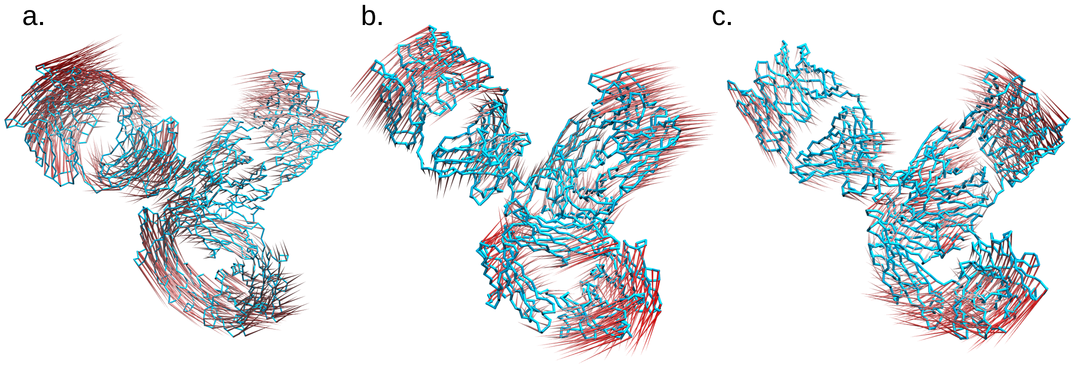


Figure S16: Porcupine representations of modes 1 (**a**), 2 (**b**) and 3 (**c**).

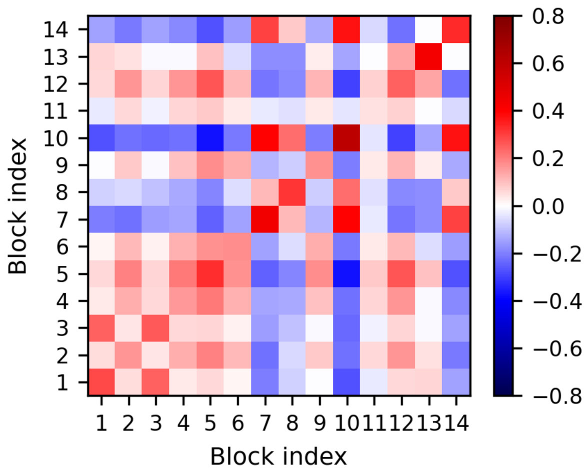


Figure S17: Collinearity of the structural domains in the first three modes.

#### Cluster $4_A$

Fab1 tends to assume a bent conformation, similar to the one in cluster $0_H$; the driving force is the concerted motion of CL in chain A, and CH1, hinge and CH2 of chain B. However, the simultaneous rotation of both CH3 domains in Fc prevents the contact of the two domains, and keeps the overall structure open. The rotations of hinge 2 and CH1 in Fab2 are also responsible; they are collinear with the corresponding domains in Fab1, but anticorrelated to the variable region of Fab2.

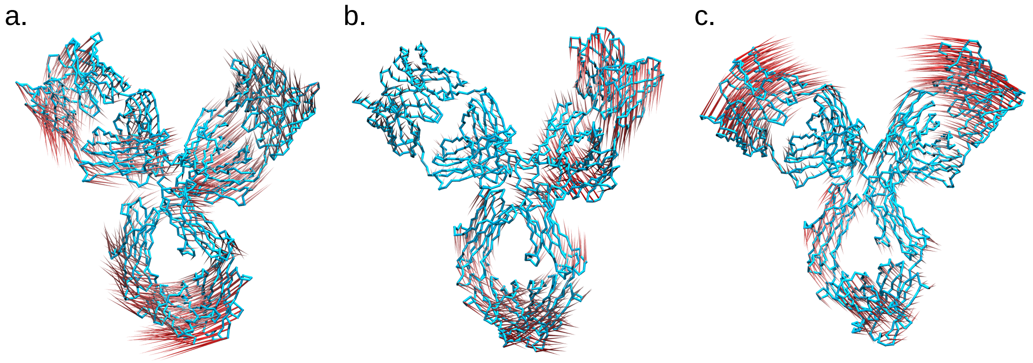


Figure S18: Porcupine representations of modes 1 (**a**), 2 (**b**) and 3 (**c**).

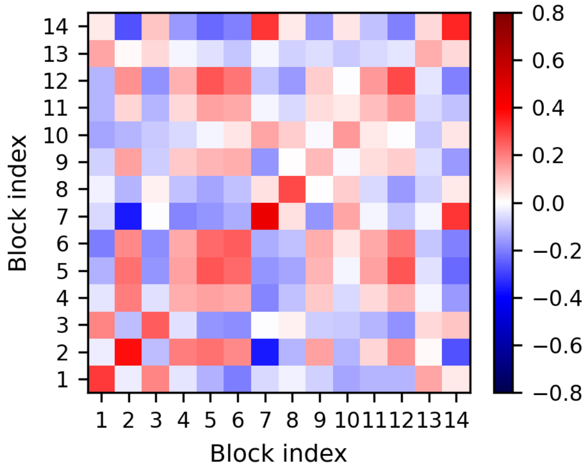


Figure S19: Collinearity of the structural domains in the first three modes.

### Cluster $5_A$

Fab1 rotates up to the point of laying perpendicular with respect to the main axis of Fc, giving to the immunoglobulin an overall T shape. This is facilitated by the simultaneous antiparallel motion of Fab2 with respect to hinge and CH2 of chain A.

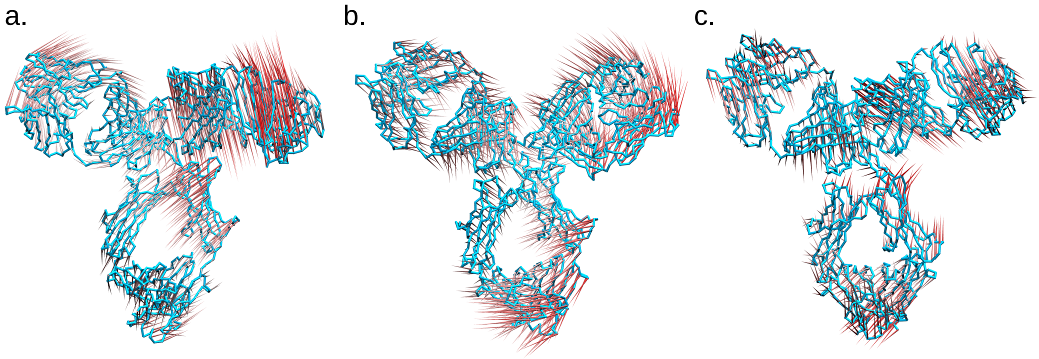


Figure S20: Porcupine representations of modes 1 (**a**), 2 (**b**) and 3 (**c**).

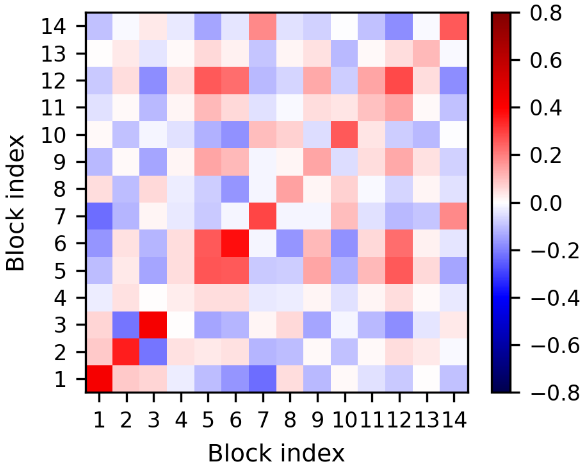


Figure S21: Collinearity of the structural domains in the first three modes.

### Cluster $0_H$

Fab1 moves in same direction of hinge, and opposite to variable domains of Fab2, in a scissor-like movement (bending). As in other clusters, a correlation between the variable region of Fab2 and the CH3 domains is observed. These movements are allowed by torsion of the lower hinge 2, and of the loop between VH and CH1 of Fab2. The motion giving rise to the peculiar conformation of Fab1 is described by mode 3; it originates from the torsion in the upper hinge of chain 1 (residues $LYS^{439}$-$TYR^{440}$-$GLY^{441}$). The conformation with bent Fab1 is then stabilized by side-chain/side-chain interactions between Fab1 and a turn in CH2 domain.

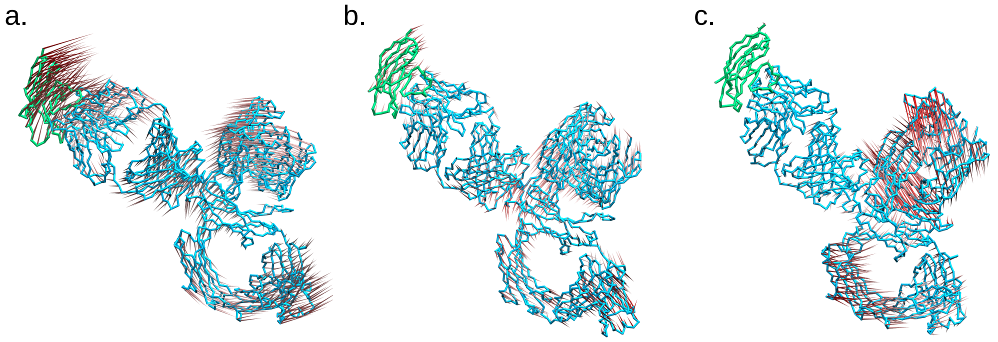


Figure S22: Porcupine representations of modes 1 (**a**), 2 (**b**) and 3 (**c**). In green is the antigen, PD-1.

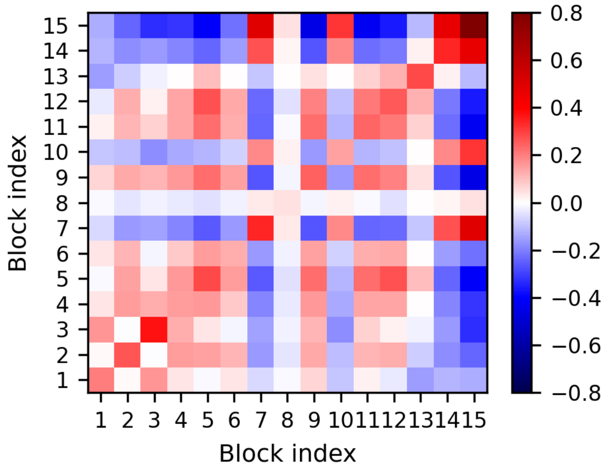


Figure S23: Collinearity of the structural domains in the first three modes.

#### Cluster $1_H$

A strongly parallel movement of the whole chain B is observed (with the usual exception of the CH3 domain). Scissoring of Fab2/Fc, until Fab2 is in contact with CH2 of chain B (many non-bonded interactions are detected, see below). In mode 2, wagging of the two Fabs and the Fc. In mode 3, a twist of the two Fabs along their major axis is observed.

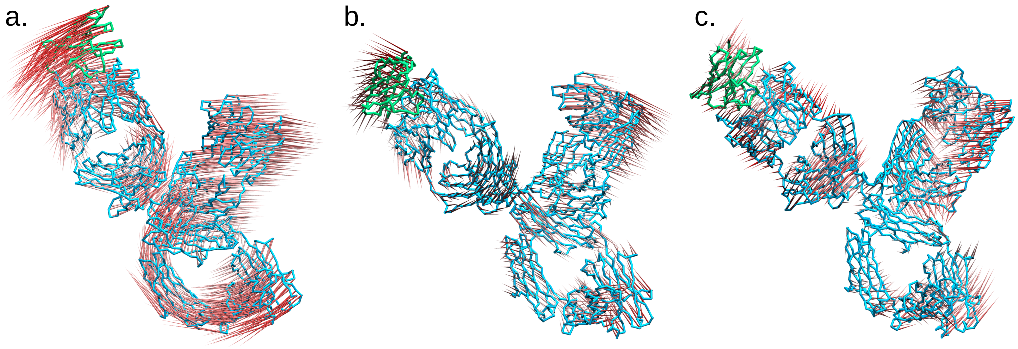


Figure S24: Porcupine representations of modes 1 (**a**), 2 (**b**) and 3 (**c**). In green is the antigen, PD-1.

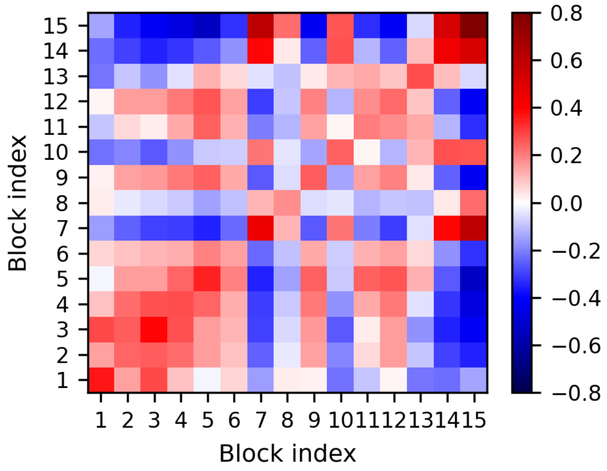


Figure S25: Collinearity of the structural domains in the first three modes.

#### Cluster $2_H$

Similar to the case of cluster $3_H$, but with a little role of Fab2 in determining the overall shape of the molecule. Wagging of Fab1 and Fc, with CL and CH2 getting close and far apart, oscillating between open and compact conformations. In mode 2 and 3, instead, Fab1 and Fc maintain a significant contact area, while Fab2 is twisted (the hinge of the motion is located on the loops connecting the variable to the constant domains).

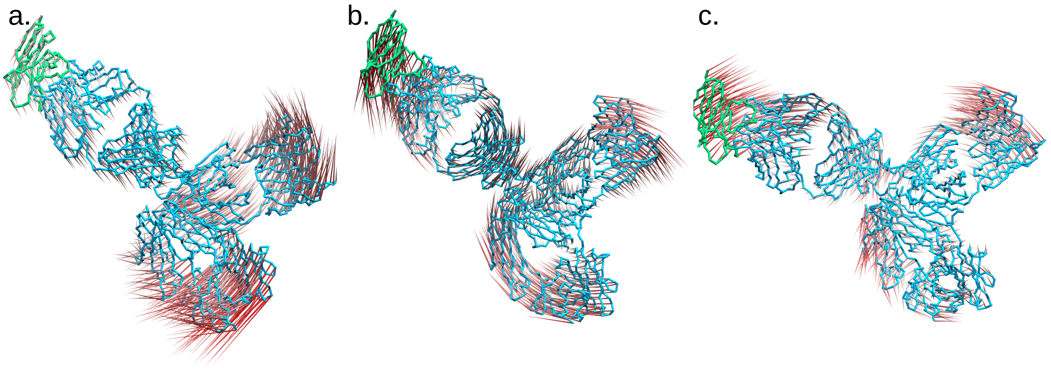


Figure S26: Porcupine representations of modes 1 (**a**), 2 (**b**) and 3 (**c**). In green is the antigen, PD-1.

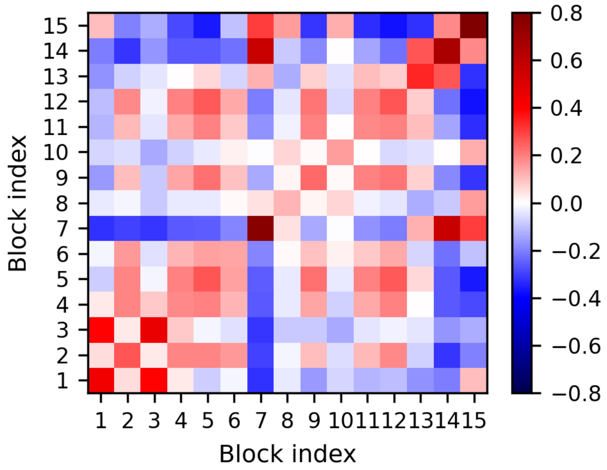


Figure S27: Collinearity of the structural domains in the first three modes.

#### Cluster $3_H$

A bending movement of the two Fabs is observed, accompanied by a bending of the two CH2 domains. An antiparallel movement of VH and hinge/VL of chain B is observed. A slight twist of Fab1 takes place in a manner similar to cluster $0_H$.

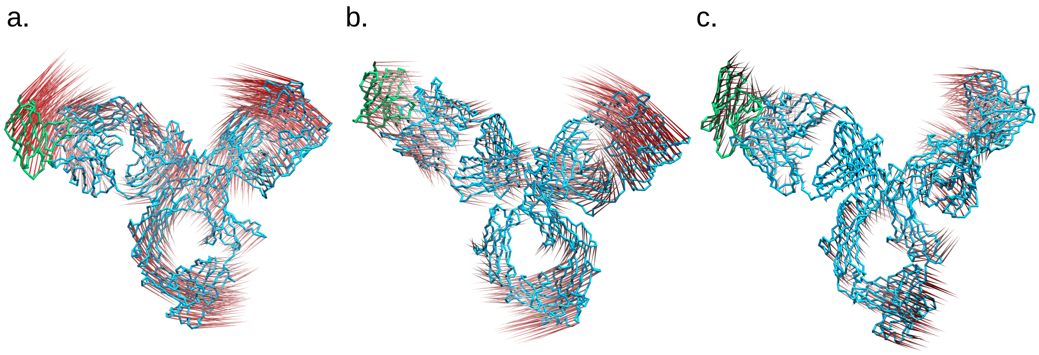


Figure S28: Porcupine representations of modes 1 (**a**), 2 (**b**) and 3 (**c**). In green is the antigen, PD-1.

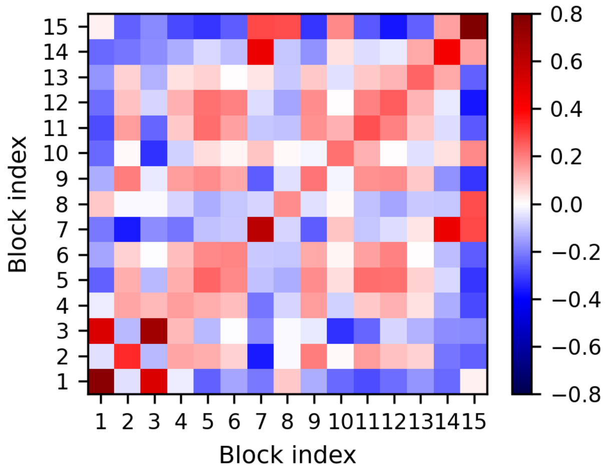


Figure S29: Collinearity of the structural domains in the first three modes.

Table S3: Residue-residue contacts characterizing the conformational clusters of the apo system. Interactions have been identified with PyContact [19]

| Cluster ID | Fab1-Fc | Fab2-Fc | Fab1-Fab2 |
|---|---|---|---|
| $0_A$ | THR$^{356}$-ILE$^{550}$,<br>VAL$^{114}$-ILE$^{595}$,<br>THR$^{113}$-MET$^{646}$,<br>THR$^{113}$-ILE$^{595}$,<br>SER$^{357}$-GLU$^{551}$,<br>THR$^{356}$-GLU$^{551}$,<br>VAL$^{114}$-ASP$^{594}$,<br>THR$^{113}$-HIS$^{647}$,<br>THR$^{113}$-ASP$^{594}$,<br>LYS$^{111}$-GLU$^{598}$,<br>VAL$^{114}$-ALA$^{596}$,<br>ASN$^{156}$-THR$^{1215}$,<br>THR$^{113}$-GLU$^{648}$,<br>THR$^{113}$-HIS$^{653}$,<br>THR$^{113}$-ALA$^{596}$,<br>ARG$^{112}$-MET$^{646}$,<br>ASN$^{156}$-LYS$^{1214}$ | No contacts | No contacts |
| $1_A$ | SER$^{357}$-THR$^{553}$,<br>SER$^{359}$-ILE$^{554}$,<br>SER$^{357}$-LYS$^{552}$,<br>SER$^{357}$-GLU$^{551}$,<br>SER$^{359}$-THR$^{553}$,<br>GLU$^{358}$-THR$^{553}$,<br>LEU$^{205}$-LYS$^{558}$,<br>VAL$^{209}$-LYS$^{558}$,<br>VAL$^{209}$-ALA$^{557}$,<br>SER$^{359}$-LYS$^{552}$,<br>SER$^{207}$-LYS$^{558}$,<br>GLU$^{358}$-SER$^{555}$,<br>THR$^{113}$-LEU$^{469}$ | No contacts | ARG$^{435}$-ASP$^{788}$,<br>LYS$^{439}$-CYS$^{880}$,<br>ARG$^{435}$-LYS$^{792}$,<br>ARG$^{435}$-LEU$^{791}$ |
| $2_A$ | THR$^{356}$-ILE$^{550}$,<br>THR$^{356}$-GLU$^{551}$,<br>SER$^{357}$-VAL$^{458}$,<br>SER$^{357}$-SER$^{457}$,<br>SER$^{357}$-GLU$^{551}$,<br>SER$^{357}$-LYS$^{552}$,<br>ARG$^{354}$-PRO$^{456}$,<br>ARG$^{354}$-SER$^{457}$,<br>GLU$^{358}$-LYS$^{552}$,<br>ARG$^{354}$-ILE$^{550}$,<br>ARG$^{354}$-LEU$^{546}$ | No contacts | No contacts |
| $3_A$ | THR$^{356}$-ILE$^{550}$,<br>SER$^{357}$-VAL$^{458}$,<br>SER$^{357}$-GLU$^{551}$,<br>SER$^{357}$-LYS$^{552}$,<br>THR$^{356}$-GLU$^{551}$,<br>ARG$^{354}$-PRO$^{456}$,<br>ARG$^{354}$-LEU$^{546}$,<br>SER$^{357}$-ILE$^{550}$ | GLU$^{879}$-ASN$^{515}$,<br>ASN$^{876}$-ASN$^{515}$,<br>GLU$^{879}$-SER$^{516}$ | LYS$^{439}$-ASP$^{788}$ |

| | | | |
|---|---|---|---|
| $4_A$ | THR$^{356}$-ASP$^{483}$, SER$^{357}$-ASP$^{483}$ | THR$^{1018}$-GLN$^{1148}$, SER$^{1074}$-LEU$^{1208}$, SER$^{1019}$-GLU$^{1149}$, ARG$^{1016}$-GLU$^{1149}$, SER$^{1019}$-ASP$^{1150}$, THR$^{1018}$-GLU$^{1149}$, SER$^{1017}$-GLU$^{1149}$ | LYS$^{439}$-ASP$^{788}$ |
| $5_A$ | No contacts | SER$^{1073}$-LEU$^{1208}$, SER$^{1074}$-LEU$^{1208}$, SER$^{1017}$-GLU$^{1149}$, THR$^{1018}$-GLU$^{1149}$, SER$^{1019}$-GLU$^{1149}$, SER$^{1019}$-PRO$^{1151}$, ARG$^{1016}$-GLU$^{1149}$, SER$^{1019}$-ASP$^{1150}$, SER$^{1021}$-LYS$^{1206}$, THR$^{1018}$-GLN$^{1148}$, GLU$^{1029}$-PRO$^{1151}$, SER$^{1021}$-GLY$^{1207}$ | ARG$^{435}$-ASP$^{788}$, CYS$^{218}$-ARG$^{1097}$, GLU$^{217}$-ARG$^{1097}$, ARG$^{435}$-LYS$^{792}$, LYS$^{439}$-ASP$^{788}$ |

Table S4: Residue-residue interactions characterizing the conformational clusters of the holo system.

| Cluster ID | Fab1-Fc | Fab2-Fc | Fab1-Fab2 |
|---|---|---|---|
| $0_H$ | $GLU^{17}$-$ARG^{473}$, $ASN^{380}$-$PHE^{514}$, $THR^{113}$-$ASP^{467}$, $PRO^{15}$-$ARG^{473}$, $THR^{113}$-$PRO^{465}$, $LYS^{417}$-$GLN^{513}$, $ARG^{112}$-$LYS^{464}$, $LEU^{384}$-$PHE^{514}$, $THR^{113}$-$LYS^{466}$, $GLU^{17}$-$ILE^{471}$, $SER^{14}$-$ARG^{473}$, $GLY^{16}$-$ILE^{471}$, $ASP^{174}$-$LYS^{464}$, $LYS^{417}$-$ASP^{483}$, $SER^{386}$-$ARG^{519}$, $ARG^{112}$-$PRO^{465}$, $ALA^{383}$-$PHE^{514}$ | $ARG^{877}$-$ASN^{515}$, $LYS^{856}$-$ASN^{515}$, $ASN^{876}$-$ASN^{515}$, $GLY^{878}$-$ASN^{515}$, $ASN^{876}$-$GLN^{486}$, $ASN^{876}$-$SER^{516}$, $LYS^{856}$-$GLU^{512}$ | $ARG^{435}$-$ASP^{788}$, $LYS^{439}$-$CYS^{880}$, $ASP^{433}$-$LYS^{849}$, $SER^{381}$-$GLU^{853}$ |
| $1_H$ | $THR^{356}$-$ILE^{550}$, $SER^{357}$-$ILE^{550}$, $SER^{359}$-$PHE^{459}$, $SER^{357}$-$LYS^{552}$, $SER^{357}$-$GLU^{551}$, $SER^{359}$-$LYS^{552}$ | $ARG^{877}$-$ASN^{515}$, $LYS^{856}$-$GLN^{486}$, $ASN^{876}$-$GLN^{486}$, $ASN^{876}$-$GLU^{487}$, $GLU^{853}$-$ASN^{515}$, $GLY^{878}$-$GLN^{486}$ | $ARG^{435}$-$ASP^{1097}$ |
| $2_H$ | $THR^{356}$-$ILE^{550}$, $SER^{357}$-$VAL^{458}$, $THR^{356}$-$GLU^{551}$, $SER^{357}$-$GLU^{551}$, $SER^{357}$-$LYS^{552}$, $SER^{357}$-$ILE^{550}$, $SER^{357}$-$SER^{457}$ | $ARG^{1016}$-$GLU^{1149}$, $VAL^{857}$-$ASN^{515}$ | $LYS^{439}$-$CYS^{880}$ |
| $3_H$ | $SER^{357}$-$VAL^{482}$, $SER^{412}$-$THR^{517}$ | $ASN^{876}$-$GLU^{487}$, $LYS^{873}$-$GLU^{1149}$, $LYS^{856}$-$GLU^{487}$, $ARG^{877}$-$GLU^{487}$, $THR^{872}$-$GLU^{1149}$, $VAL^{871}$-$ASP^{1150}$, $GLU^{1020}$-$LYS^{1206}$, $SER^{1021}$-$LEU^{1208}$, $SER^{874}$-$GLU^{1149}$ | No contacts |

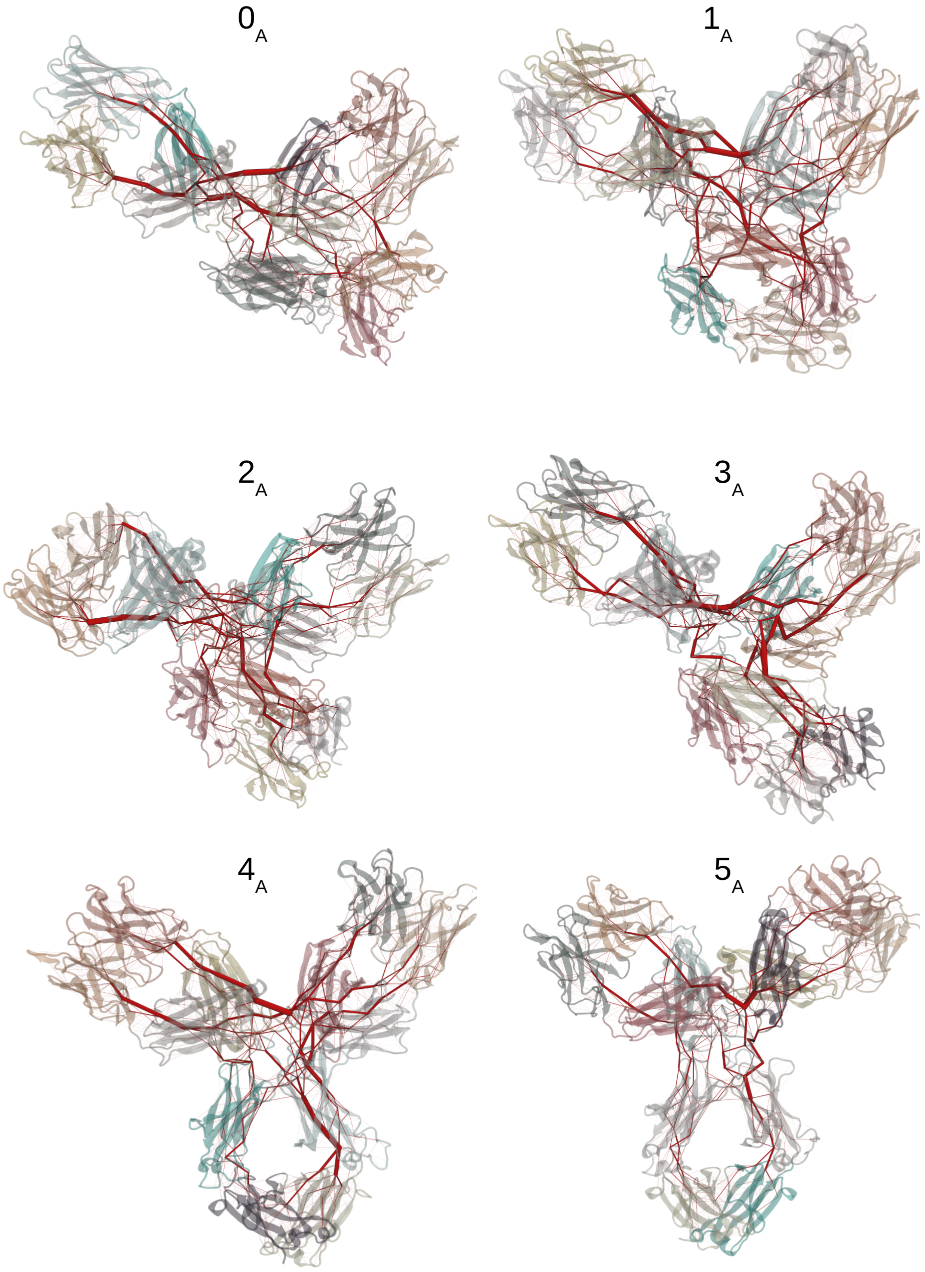


Figure S30: Edges of the apo networks are represented as red lines, with thickness proportional to the edge betweenness. The numbers indicate the cluster ID.

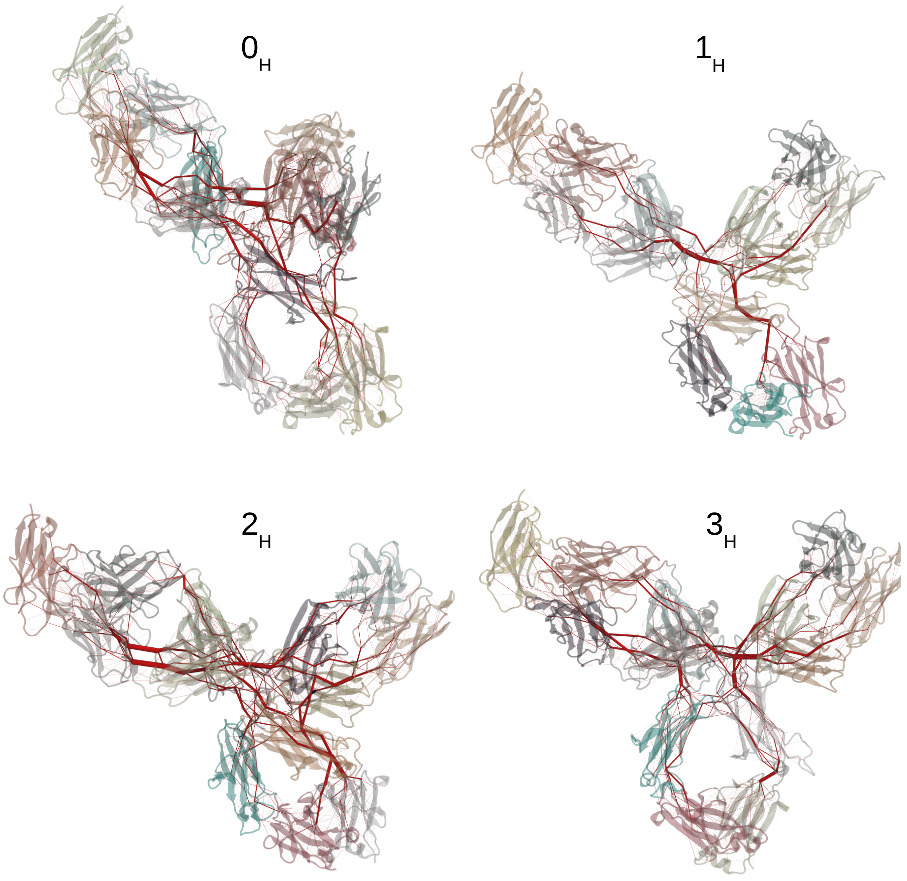


Figure S31: Edges of the holo network are represented as red lines, with thickness proportional to the edge betweenness. The numbers indicate the cluster ID.

Table S5: Edges with the highest betweenness in the apo clusters.

| Cluster ID | Residue 1 | Domain | Residue 2 | Domain |
|---|---|---|---|---|
| Cluster $0_A$ | ARG 435 | 4 | PRO 443 | 5 |
| | LEU 1076 | 11 | TYR 1102 | 12 |
| | TYR 1102 | 12 | PRO 1108 | 12 |
| | PRO 443 | 5 | GLU 1099 | 11 |
| | ASN 803 | 9 | TYR 839 | 9 |
| | CYS 352 | 4 | PRO 1108 | 12 |
| | GLU 771 | 8 | TYR 839 | 9 |
| | ASN 803 | 9 | THR 1070 | 11 |
| | LEU 1028 | 11 | SER 1064 | 11 |
| | CYS 421 | 4 | ARG 435 | 4 |
| Cluster $1_A$ | ARG 435 | 4 | ASP 788 | 9 |
| | GLN 790 | 9 | LYS 1030 | 11 |
| | LYS 1030 | 11 | TYR 1063 | 11 |
| | THR 997 | 10 | TYR 1063 | 11 |
| | CYS 421 | 4 | ARG 435 | 4 |
| | SER 412 | 4 | THR 416 | 4 |
| | LYS 558 | 6 | PRO 592 | 7 |
| | TYR 440 | 5 | PRO 442 | 5 |
| | ARG 435 | 4 | LYS 792 | 9 |
| | THR 971 | 10 | LEU 1057 | 11 |
| Cluster $2_A$ | GLU 771 | 8 | TYR 839 | 9 |
| | ASN 803 | 9 | TYR 839 | 9 |
| | ASN 803 | 9 | THR 1070 | 11 |
| | ARG 354 | 4 | ILE 550 | 6 |
| | ARG 1097 | 11 | SER 1100 | 11 |
| | SER 999 | 10 | PHE 1033 | 11 |
| | VAL 500 | 6 | LEU 616 | 7 |
| | TYR 537 | 6 | LYS 552 | 6 |
| | VAL 1085 | 11 | HIS 1087 | 11 |
| | CYS 1083 | 11 | ARG 1097 | 11 |
| Cluster $3_A$ | TYR 537 | 6 | LYS 552 | 6 |
| | SER 359 | 4 | LYS 552 | 6 |
| | SER 438 | 4 | CYS 444 | 5 |
| | PRO 442 | 5 | ALA 1012 | 11 |
| | LYS 439 | 4 | PRO 442 | 5 |
| | THR 360 | 4 | THR 408 | 4 |
| | CYS 444 | 5 | GLU 1099 | 11 |
| | THR 971 | 10 | LEU 1057 | 11 |
| | PHE 120 | 2 | THR 408 | 4 |
| | LEU 1028 | 11 | SER 1064 | 11 |
| Cluster $4_A$ | LYS 439 | 4 | ASP 788 | 9 |
| | LYS 535 | 6 | TYR 537 | 6 |
| | ASP 788 | 9 | GLN 790 | 9 |
| | ASN 803 | 9 | TYR 839 | 9 |
| | TYR 537 | 6 | CYS 539 | 6 |
| | CYS 539 | 6 | ILE 550 | 6 |
| | GLN 790 | 9 | LYS 1030 | 11 |
| | LYS 769 | 8 | TYR 839 | 9 |
| | LYS 535 | 6 | TYR 591 | 7 |
| | GLY 455 | 5 | SER 457 | 6 |

| | | | | |
|---|---|---|---|---|
| Cluster $5_A$ | GLU 437 | 4 | LYS 439 | 4 |
| | LYS 439 | 4 | ASP 788 | 9 |
| | ILE 550 | 5 | PHE 1114 | 12 |
| | ASP 788 | 9 | GLN 790 | 9 |
| | GLN 790 | 9 | LEU 1028 | 11 |
| | PRO 1112 | 12 | PHE 1114 | 12 |
| | ASN 803 | 9 | TYR 839 | 9 |
| | THR 971 | 10 | LEU 1057 | 11 |
| | PRO 1110 | 12 | PRO 1112 | 12 |
| | LEU 363 | 4 | GLU 437 | 4 |

Table S6: Edges with the highest betweenness in the holo clusters.

| Cluster ID | Residue 1 | Domain | Residue 2 | Domain |
|---|---|---|---|---|
| Cluster $0_H$ | ARG 435 | 4 | ASP 788 | 9 |
| | TYR 752 | 8 | LYS 769 | 8 |
| | TYR 419 | 4 | ARG 435 | 4 |
| | LYS 769 | 8 | ARG 808 | 9 |
| | ASP 788 | 9 | GLN 790 | 9 |
| | LYS 769 | 8 | GLU 831 | 9 |
| | GLU 512 | 6 | LYS 856 | 9 |
| | GLN 790 | 9 | LEU 1028 | 11 |
| | GLU 512 | 6 | ARG 519 | 6 |
| | LYS 535 | 6 | GLY 620 | 7 |
| Cluster $1_H$ | TYR 440 | 5 | PRO 445 | 5 |
| | PRO 445 | 5 | GLU 1099 | 11 |
| | LEU 414 | 4 | TYR 440 | 5 |
| | TYR 537 | 6 | LYS 552 | 6 |
| | LYS 535 | 6 | TYR 537 | 6 |
| | LEU 1025 | 11 | GLU 1099 | 11 |
| | ASN 141 | 2 | TYR 177 | 2 |
| | SER 359 | 4 | LYS 552 | 6 |
| | LYS 535 | 6 | LEU 616 | 7 |
| | PHE 120 | 2 | THR 360 | 4 |
| Cluster $2_H$ | ASN 803 | 9 | TYR 839 | 9 |
| | GLU 771 | 8 | TYR 839 | 9 |
| | LYS 769 | 8 | GLU 831 | 9 |
| | LYS 535 | 6 | TYR 537 | 6 |
| | LYS 535 | 6 | TYR 591 | 7 |
| | LYS 439 | 4 | CYS 880 | 9 |
| | ASN 141 | 2 | TYR 177 | 2 |
| | THR 830 | 9 | GLU 831 | 9 |
| | PRO 785 | 9 | CYS 880 | 9 |
| | TYR 752 | 8 | LYS 769 | 8 |
| Cluster $3_H$ | LEU 414 | 4 | TYR 440 | 5 |
| | TYR 440 | 5 | PRO 445 | 5 |
| | LEU 453 | 5 | GLY 455 | 5 |
| | PRO 445 | 5 | GLU 1099 | 11 |
| | TYR 752 | 8 | LYS 769 | 8 |
| | ASN 533 | 6 | TYR 591 | 7 |
| | LYS 873 | 9 | GLU 1149 | 13 |
| | LEU 363 | 4 | LEU 414 | 4 |
| | ASN 141 | 2 | TYR 177 | 2 |
| | ASN 803 | 9 | TYR 839 | 9 |

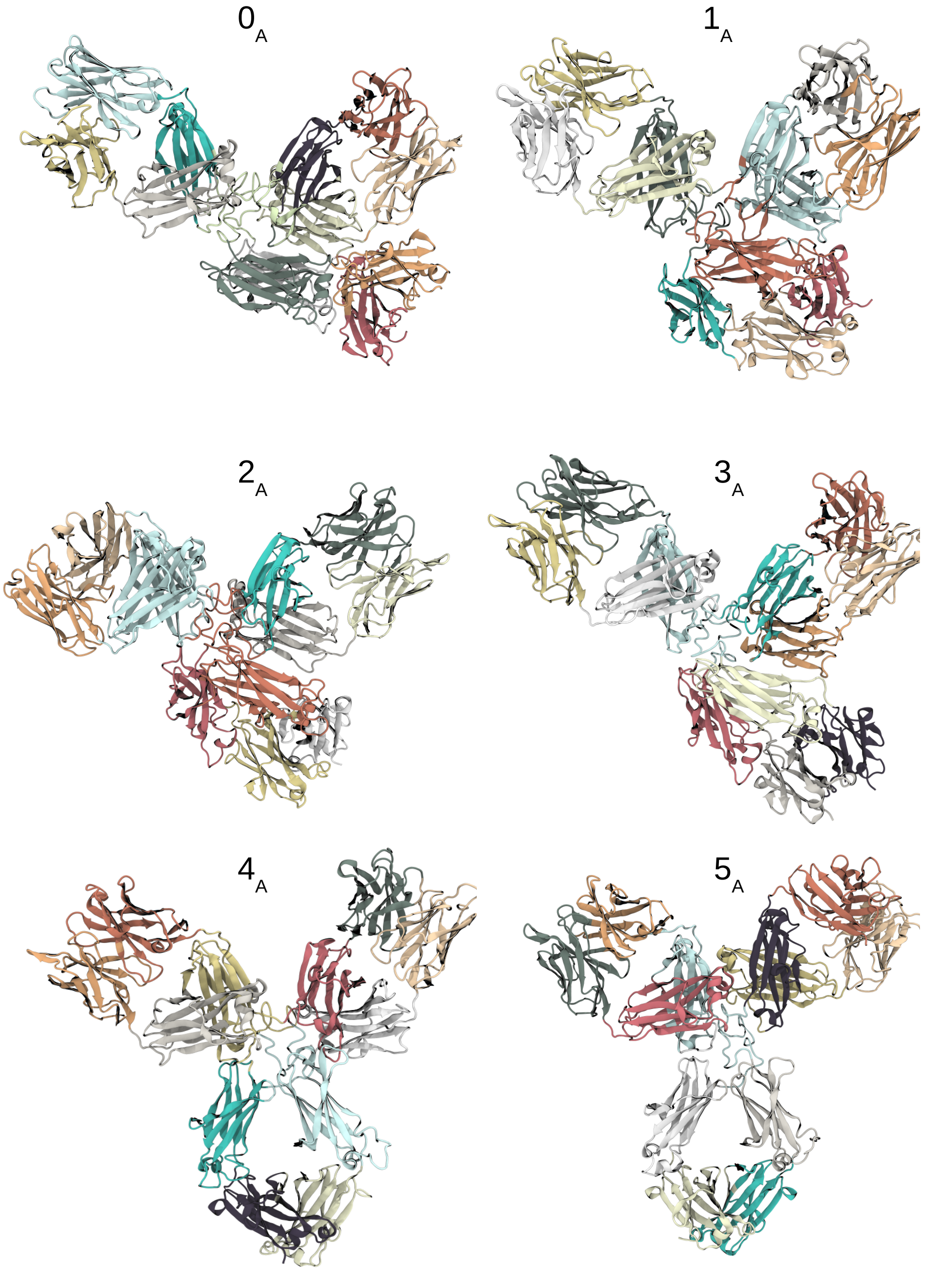


Figure S32: Community network repartitions for the apo system. The numbers indicate the cluster ID.

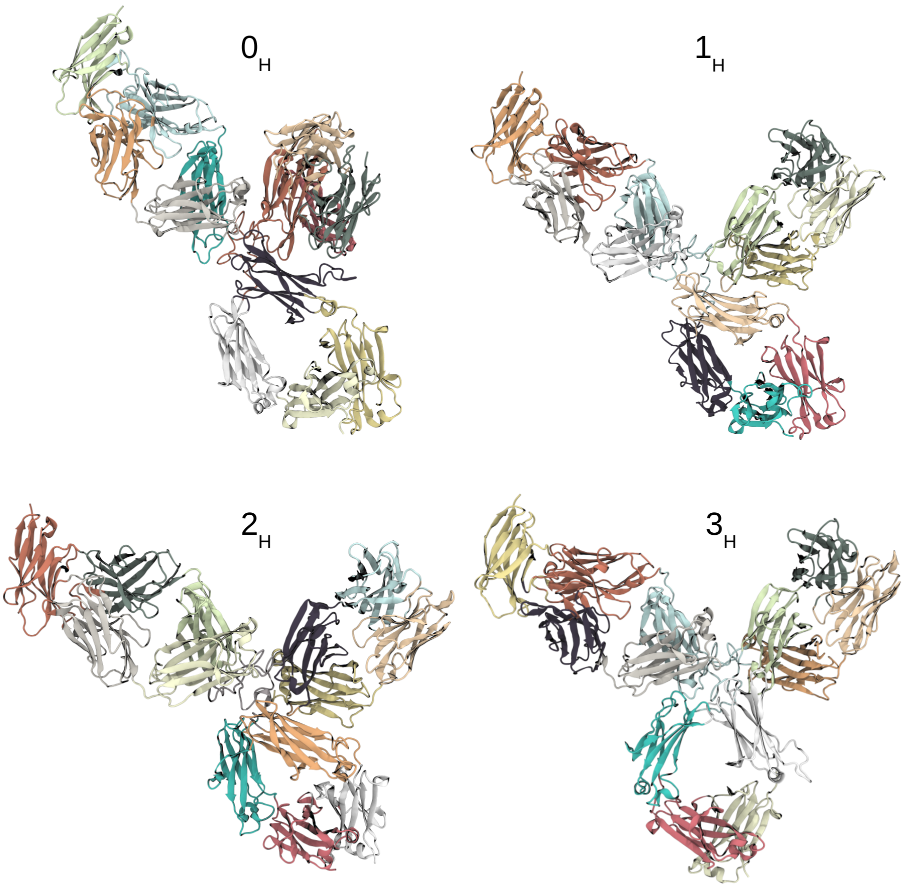


Figure S33: Community network repartitions for the holo system. The numbers indicate the cluster ID.

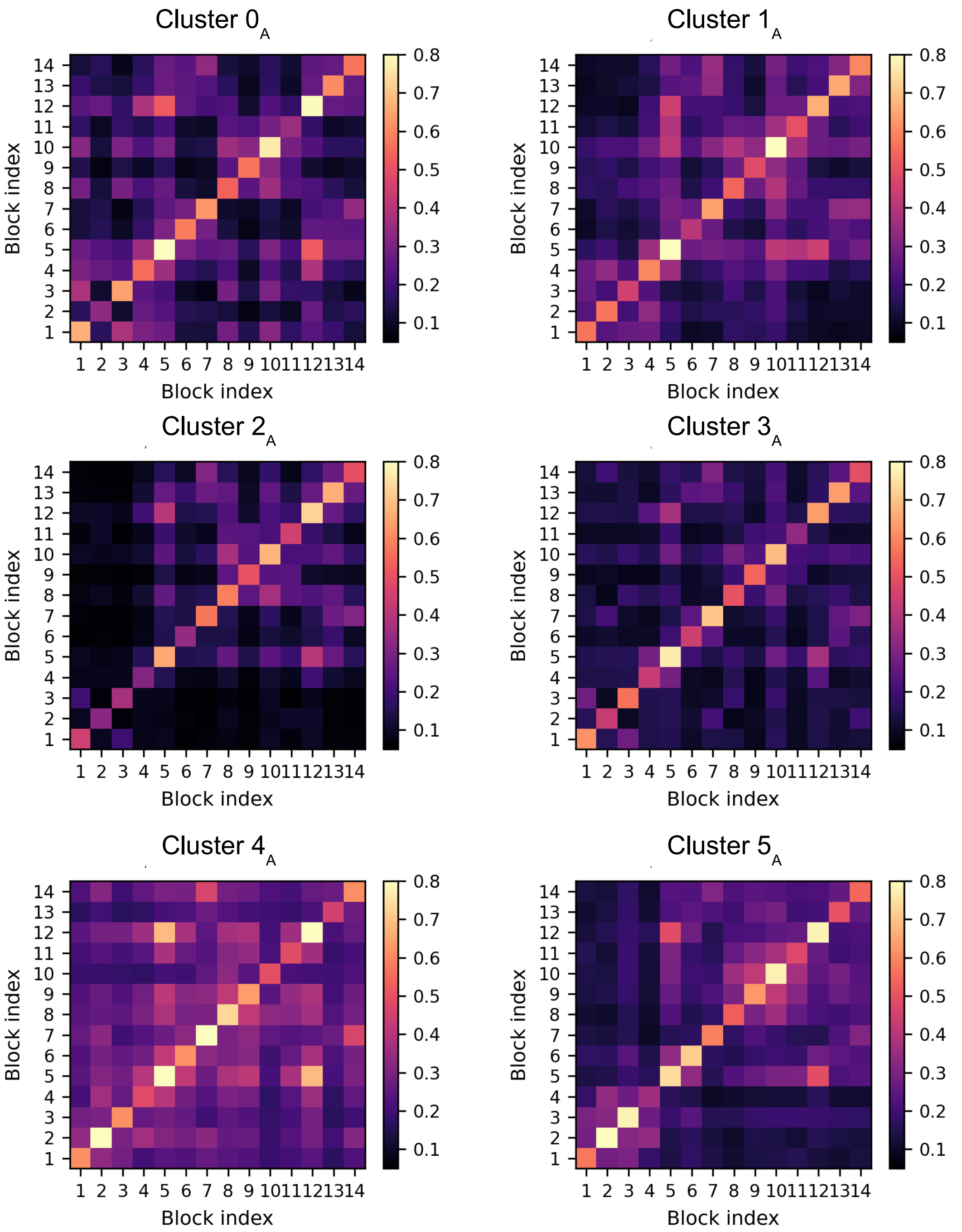


Figure S34: Mutual information averaged within each structural domain of pembrolizumab in the apo state, for each conformational cluster.

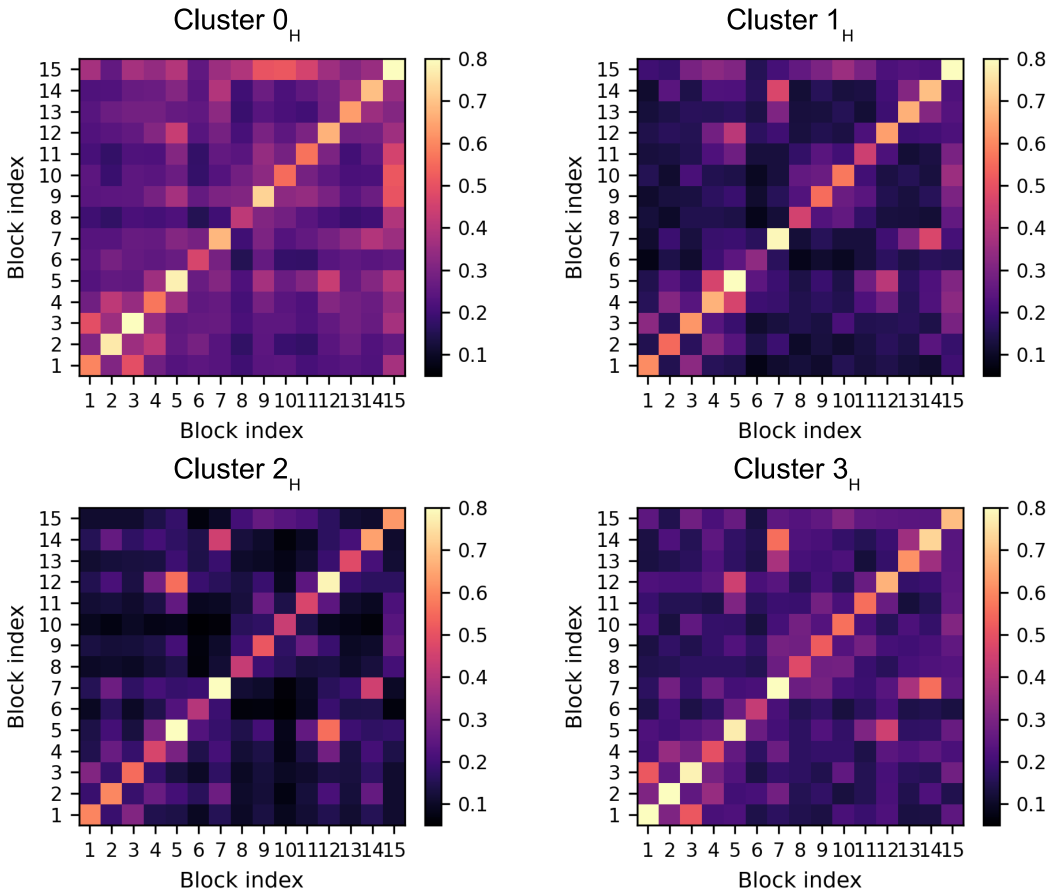


Figure S35: Mutual information averaged within each structural domain of pembrolizumab in the holo state, for each conformational cluster.

### S2.2 The hinge region

Hinge 1, namely the hinge segment of chain B, shows higher correlations than hinge 2 with the rest of the protein, in all the apo and holo clusters. These correlations are particularly strong with the domains of Fab2. Network analysis shows that, in the apo case, a particularly favoured edge is the one between $ASP^{788}$ and $LYS^{439}$; the former is part of Fab2, and the latter is connecting Fab1 and the hinge segment. Atomistic-level inspection shows the presence of a persistent salt bridge between the two residues in clusters $3_A$, $4_A$ and $5_A$.
Edge betweenness highlights the importance of hinge proline residues for the compact apo conformations (cluster $0_A$). In particular, $PRO^{443}$ in hinge 1 and $PRO^{1108}$ in hinge 2 are key residues for inter-Fabs communication. The former is part of the highly central path $ARG^{435}$-$PRO^{443}$-$GLU^{1099}$, while the latter (which corresponds to the residue mutated from serine, in order to avoid Fab-arm exchange [20]) is part of $CYS^{352}$-$PRO^{1108}$-$TYR^{1102}$-$LEU^{1076}$. This last path is observed also in other compact conformations, those of cluster $2_A$; however, in this case, the network involving hinge residues is fragmented in a large number of paths with low centrality.
Prolines play a central role also in cluster $1_A$, where $PRO^{442}$ in hinge 1 is highly correlated to $PRO^{1010}$ on Fab2. $PRO^{442}$ is also part of a highly central path connecting hinge 1 with Fab1 through $TYR^{440}$ and $THR^{416}$.
The same $PRO^{442}$ residue is involved in a highly central path in cluster $3_A$, where it connects $LYS^{439}$ in Fab1 to $ALA^{1012}$ in Fab2. In this cluster, the path involving residues $SER^{438}$-$LYS^{444}$-$GLU^{1099}$ is also highly relevant.

In the holo case we observe that, also in the most open conformations (cluster $3_H$), the hinge region still plays an important role for information transfer, in particular when compared to the apo case. Again, prolines appear as key residues. In cluster $0_H$, there are no significant interactions between the hinge residues and the nearby domains. In all clusters, none of the residues of hinge 2 is involved in high-centrality contacts. Several paths with highest betweenness include instead hinge 1 residues. In cluster $2_H$, pathways with high betweenness include: $PRO^{1013}$-$SER^{1015}$-$PHE^{451}$-$GLY^{455}$, $CYS^{879}$-$LYS^{439}$ and $PRO^{445}$-$CYS^{1105}$. In clusters $1_H$ and $3_H$ there is one path with high centrality crossing the hinge, namely $LEU^{414}$-$TYR^{440}$-$PRO^{445}$-$GLU^{1099}$.

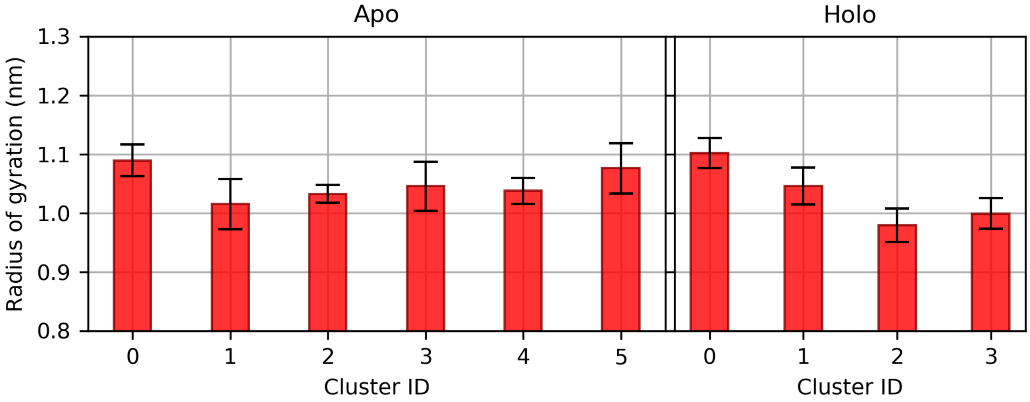


Figure S36: Average radii of gyration of the hinge region, in the apo and holo clusters.

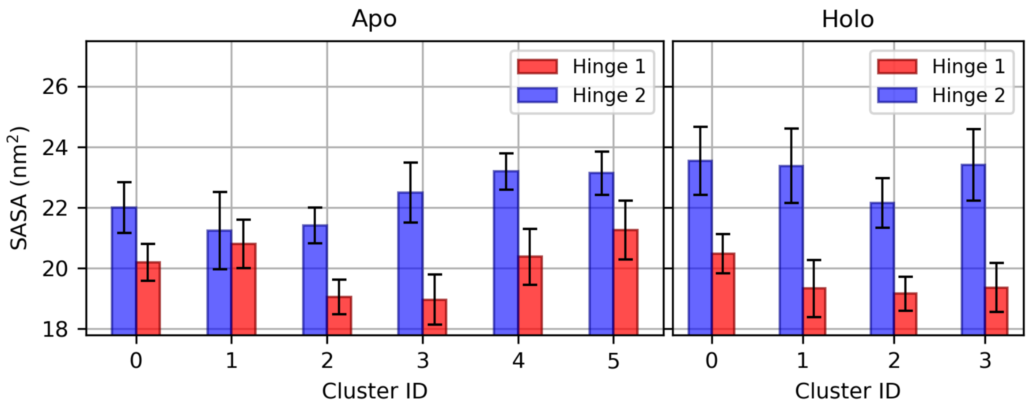


Figure S37: Average solvent accessible surface area (SASA) of hinge segments in chain B (hinge 1) and chain D (hinge 2), in the apo and holo clusters. The values refer to the hinge surface alone, computed without taking into account hindrance from nearby domains. The two segments appear highly asymmetrical; in particular, the smaller SASA for hinge 1 is indicative of a more compact, less accessible conformation.

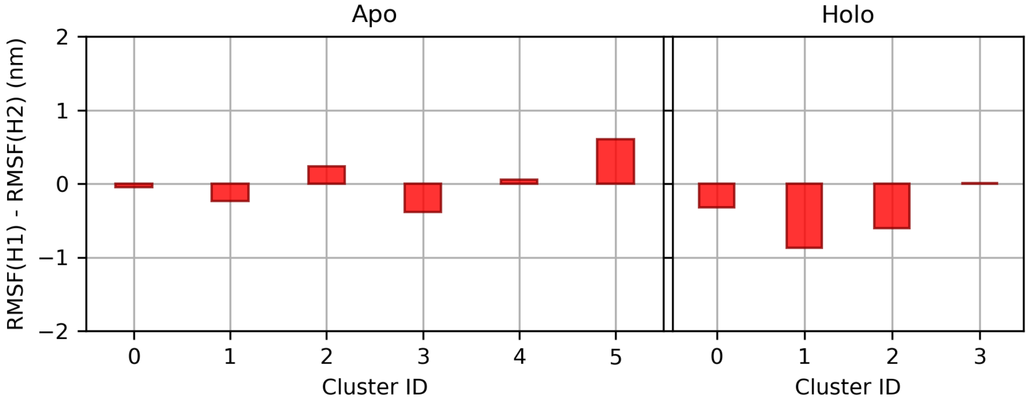


Figure S38: Difference between the total $C^{\alpha}$ RMSF of the two hinge chains, in the clusters of apo and holo states. In the holo case, hinge 1 appears overall less flexible than hinge 2, while in the apo systems this is true only for 2 of the 5 conformational clusters.

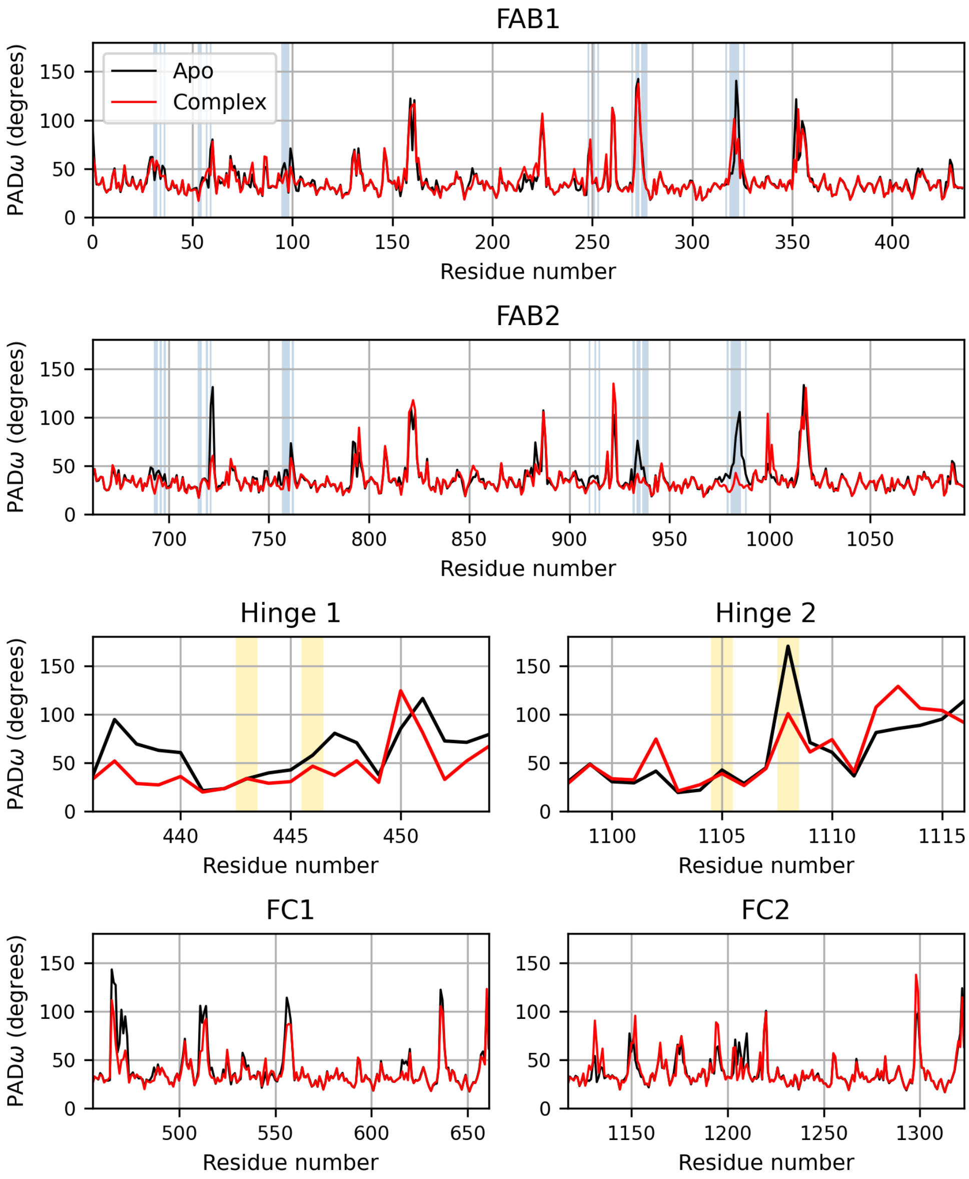


Figure S39: Comparison between the PAD$_\omega$ parameter in the apo and holo antibody. PAD$_\omega$ measures of backbone torsional plasticity through the spreading of $\phi$ and $\psi$ dihedral angles. The blue-shaded areas correspond to the residues involved in the binding with PD1; as expected, the bound antigen leads to a decreased flexibility in the paratope of Fab2. The yellow-shaded areas correspond to the cysteine residues forming inter-chain disulfide bonds in the hinge.

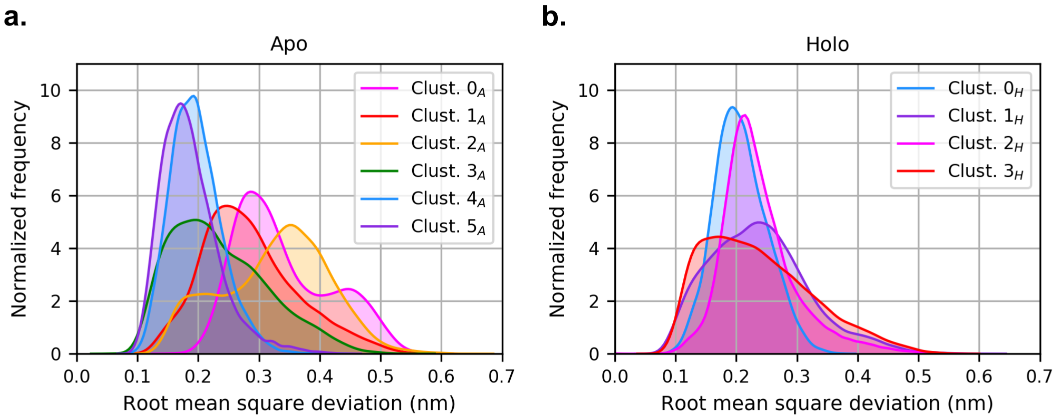


Figure S40: Plots of the RMSD of binding site residues ($C^\alpha$) in the conformational clusters of the apo (**a**) and holo (**b**) systems, with respect to the crystallographic structure of the Fab/PD-1 complex. The deviations are calculated after fitting the trajectories on the variable region (Fab2) of the antibody. In the holo case, clusters present a more uniform behaviour with respect to the apo system.

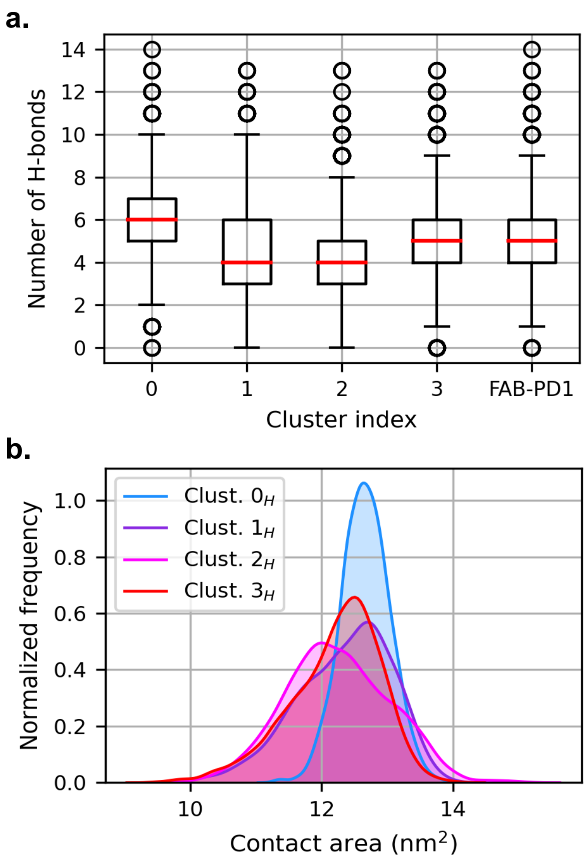


Figure S41: **a.** Comparison between the number of hydrogen bonds between the CDR and the PD1 in the four clusters of the bound antibody, and in the simulation of the antigen with the Fab alone. **b.** Comparison of the contact area distributions between the Fv and the PD1 in the four clusters of the bound antibody.

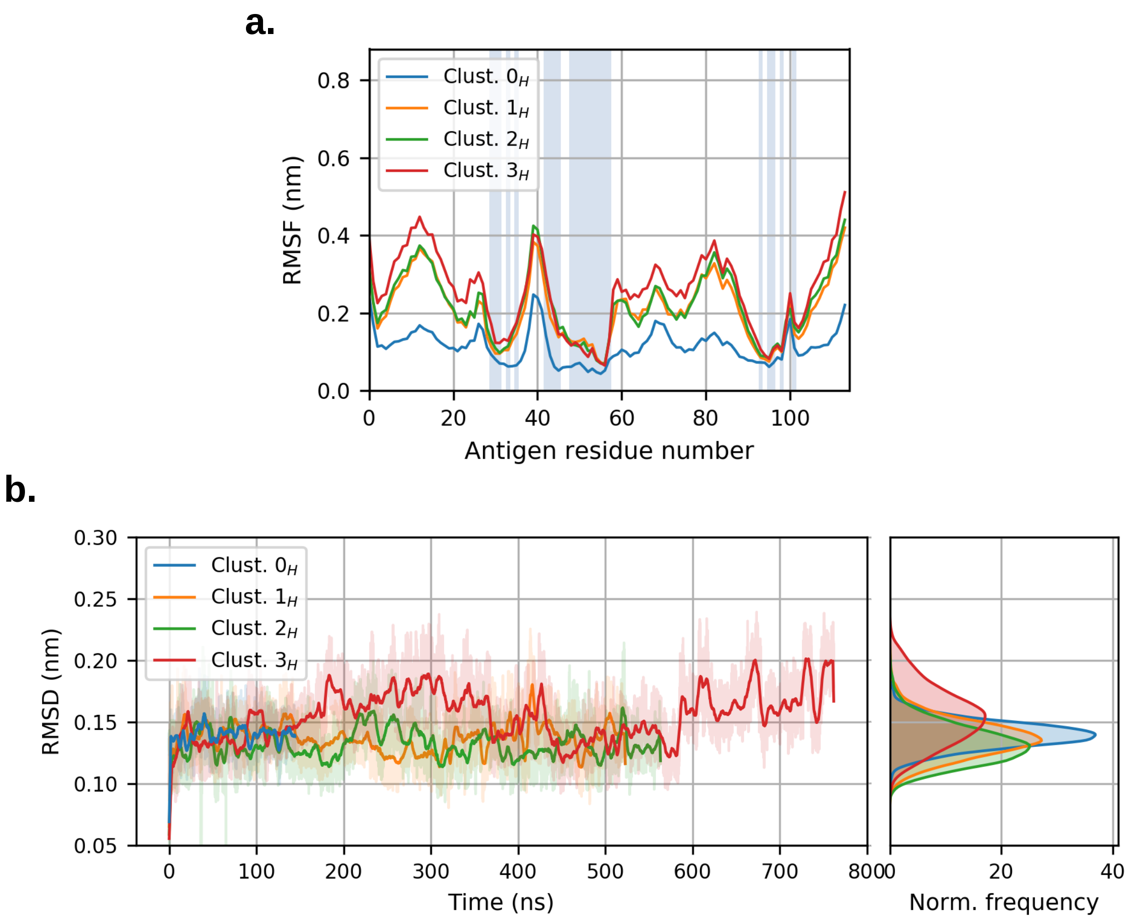


Figure S42: **a.** Root-mean-square fluctuations (RMSF) of $C^{\alpha}$ atoms of antigen PD-1, for each of the four clusters. The shaded areas correspond to residues of the epitope. **b.** Root-mean-square deviation (RMSD) of $C^{\alpha}$ atoms of PD-1 with respect to the starting conformation, and corresponding distributions.

Table S7: Enthalpic contributions to the antibody-PD1 binding energy, with respect to the value from the simulation of the Fab alone in complex with PD-1. Given the physical approximations, the results should be interpreted at a qualitative level.

| Cluster ID | $\Delta\Delta H$ (Kcal/mol) |
|---|---|
| $0_H$ | $-24.1 \pm 0.7$ |
| $1_H$ | $-16.0 \pm 0.7$ |
| $2_H$ | $-17.5 \pm 0.8$ |
| $3_H$ | $-22.0 \pm 0.7$ |

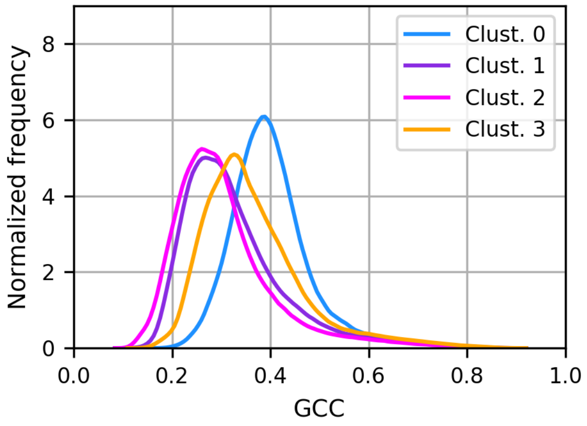


Figure S43: Distributions of the generalized correlation coefficient (normalized mutual information) within the antibody in the holo state, for each of the four clusters. The trend roughly follows the strength of the antibody-antigen binding.

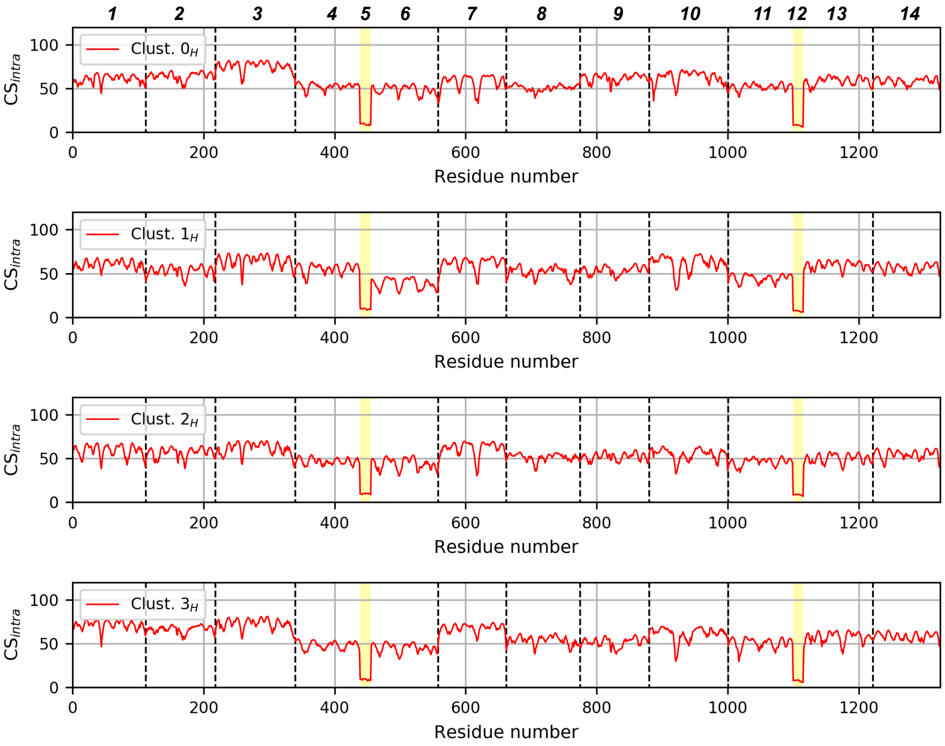


Figure S44: Intra-domain correlation score for each residue of the antibody in the holo state. The dashed vertical lines represent the boundaries of the structural domains, while the yellow-shaded areas the hinge regions. The indices on top refer to the numbering of the structural domains as in S3.

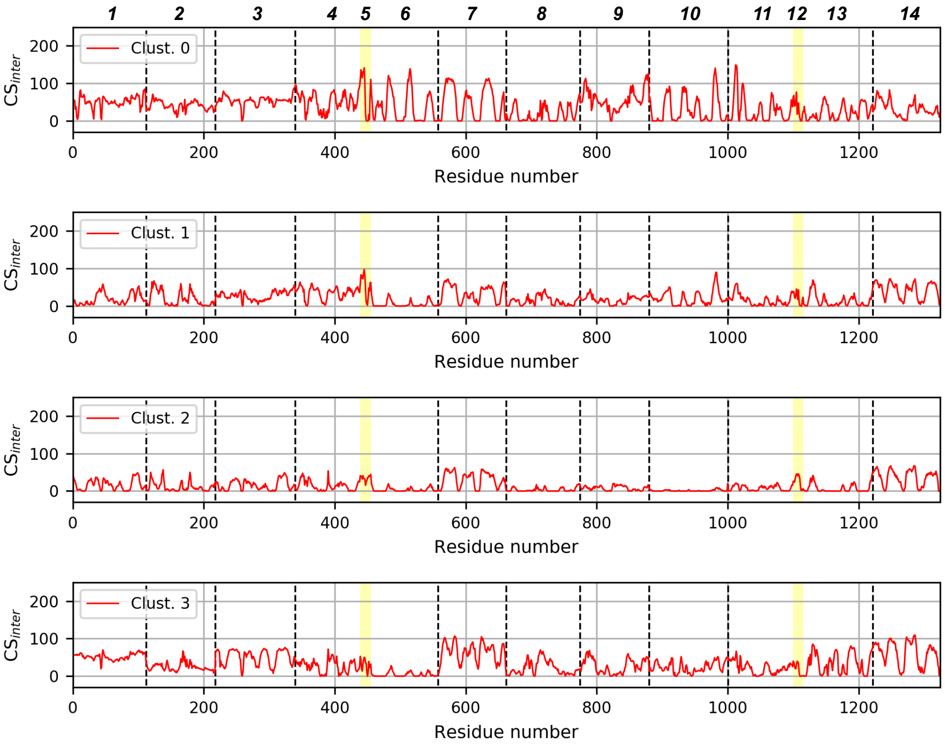


Figure S45: Inter-domain correlation coefficients for each residue of the bound antibody. The dashed vertical lines represent the boundaries of the structural domains, while the yellow-shaded areas the hinge regions. In order to facilitate the comparison, $CS_{inter}$ values have been plotted after filtering low-value correlations (GCC<0.4). The indices on top refer to the numbering of the structural domains as in S3.

| Block 15 | Cluster $0_H$ | Cluster $1_H$ | Cluster $2_H$ | Cluster $3_H$ |
|---|---|---|---|---|
| Block 1 | 0.36 | 0.19 | 0.11 | 0.24 |
| Block 2 | 0.26 | 0.19 | 0.11 | 0.15 |
| Block 3 | 0.37 | 0.30 | 0.11 | 0.28 |
| Block 4 | 0.33 | 0.32 | 0.13 | 0.21 |
| Block 5 | 0.40 | 0.31 | 0.18 | 0.27 |
| Block 6 | 0.26 | 0.15 | 0.07 | 0.13 |
| Block 7 | 0.35 | 0.20 | 0.1 | 0.25 |
| Block 8 | 0.38 | 0.27 | 0.21 | 0.24 |
| Block 9 | 0.52 | 0.31 | 0.27 | 0.26 |
| Block 10 | 0.51 | 0.36 | 0.24 | 0.31 |
| Block 11 | 0.45 | 0.31 | 0.22 | 0.26 |
| Block 12 | 0.36 | 0.22 | 0.17 | 0.25 |
| Block 13 | 0.30 | 0.24 | 0.12 | 0.25 |
| Block 14 | 0.35 | 0.22 | 0.11 | 0.24 |
| Block 15 | 1.07 | 0.78 | 0.60 | 0.66 |

Figure S46: Mutual information values for the holo cluster, between antigen (block 15) and all the structural domains of the antibody. The coloured blocks represent the highest values of each pair: in red the values grater than 0.5, in light blue the only 2 inter-domain correlations that are comparable between cluster $0_H$ and any of the other clusters ($1_H$).

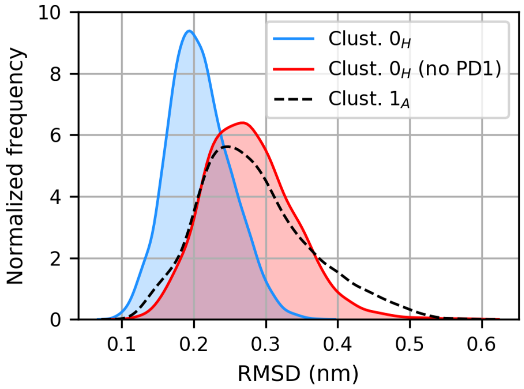


Figure S47: Distributions of the binding site RMSD values, in the holo cluster with the tightest binding and the strongest inter-protein correlations (namely cluster $0_H$); in the structurally most similar apo cluster (cluster $1_A$); and in the simulations started from the representative structure of $0_H$ after removal of the antigen.
